# Oncogenic fusions induce an extensive cancer-restricted cryptic proteome in Ewing sarcoma

**DOI:** 10.64898/2026.09.16.751792

**Authors:** Luuk A. Broeils, Jill Pilet, Mariia K. Koshkina, Jing Li, Sandrine Grossetete, Emma G. A. Westerink, Orson Jay, Lotta Smit, Ting Luo, Erkan Narmanli, Ana Pinheiro Lopes, Florian H. Geyer, Sem A. G. Engels, Clémence Henon, Karim Aljakouch, Jeroen Krijgsveld, Ismail Jamail, Sylvain Baulande, Sonia Lameiras, Martha J. Carreño-Gonzalez, Georg W. Omlor, Jonathan M. Mudge, Valerie U. Nguyen, Michal I. Swirski, Hakon Tjeldnes, Eivind Valen, Burkhard Lehner, Uta Dirksen, Wolfgang Faigle, Ana I. Lalanne, Olivier Lantz, Karine Laud-Duval, Christina Michail, Michael VanInsberghe, Alexander van Oudenaarden, Nina Hahnen, Nadine Gmelin, Johannes H. M. Merks, Ferhat Alkan, Joshua J. Waterfall, Thomas G. P. Grünewald, Olivier Delattre, Sebastiaan van Heesch

**Affiliations:** Princess Máxima Center for Pediatric Oncology, Utrecht, The Netherlands; Oncode Institute, Utrecht, The Netherlands; INSERM U1330, CNRS EMR 8001, Diversity and Plasticity of Childhood Sarcoma Lab, PSL Research University, SIREDO Oncology Center, Institut Curie Research Center, Paris, France; INSERM U1330, Integrative Functional Genomics of Cancer Lab, PSL Research University, Department of Translational Research, PSL Research University, Institut Curie Research Center, Paris, France; Division of Translational Pediatric Sarcoma Research, German Cancer Research Center (DKFZ), German Cancer Consortium (DKTK), Heidelberg, Germany; Hopp Children’s Cancer Center Heidelberg (KiTZ), Heidelberg, Germany; National Center for Tumor Diseases (NCT), NCT Heidelberg, a partnership between DKFZ and Heidelberg University Hospital, Germany; Faculty of Medicine, Heidelberg University, Heidelberg, Germany; Division of Proteomics of Stem Cell and Cancer, German Cancer Research Center (DKFZ), Heidelberg, Germany; ICGex Next-Generation Sequencing Platform, PSL Research University, Institut Curie Research Center, 75005 Paris, France; CUBIC Bioinformatics Core Facility, PSL Research University, Institut Curie Research Center, 75005 Paris, France; Laboratory of Clinical immunology, Department of Diagnostic and Theranostic Medicine, Institut Curie, Paris, France; Centre d’investigation Clinique Inserm 2501, Institut Curie, Paris, France; INSERM U932, Immune responses and cancer Lab, PSL Research University, Institut Curie Research Center, Paris, France; Hubrecht Institute-KNAW (Royal Netherlands Academy of Arts and Sciences), Utrecht, the Netherlands; Institute of Pathology, Heidelberg University Hospital, Heidelberg, Germany; Department of Orthopaedics, Heidelberg University Hospital, Heidelberg, Germany; Center for Orthopedics and Joint Replacement, Marienhaus Hospital, 66606 St. Wendel, Germany; European Molecular Biology Laboratory, European, Bioinformatics Institute, Wellcome Genome Campus, Hinxton, Cambridge, UK; Department of Biosciences, University of Oslo, 0316 Oslo, Norway; Pediatrics III, University Hospital Essen, West German Cancer Center, Essen, Germany; National Center for Tumor Diseases (NCT) Site Essen, German German Cancer Consortium (DKTK), Essen, Germany; University Medical Center Utrecht, Utrecht, The Netherlands; Division of Imaging and Oncology, University Medical Center Utrecht, Utrecht University, Utrecht, the Netherlands

**Author notes:** shared senior authorship and corresponding authors. equally contributed (Luuk A. Broeils, Jill Pilet, Mariia K. Koshkina).

**Keywords:** Ewing sarcoma, neogenes, dark proteome, antigen discovery, non-canonical translons, Ribo-seq, long-read transcriptomes, *de novo* gene evolution, cancer vaccines, immunotherapy

## Abstract

How tumors generate a cryptic “dark” proteome absent from healthy tissues, and whether it can be reversibly activated, remains unclear. In Ewing sarcoma, all tumors are driven by EWSR1::ETS fusions, making it a tractable model to study dark proteome activation. Integrating matched transcriptomes, translatomes and proteomes from 48 patient tumors with long-read RNA sequencing, single-cell Ribo-seq, proteomics and immunopeptidomics in cell line models, we show that EWSR1::FLI1 acts as a reversible switch recurrently inducing hundreds of cancer-specific neoproteins. Many arise from canonical coding regions via intragenic transcription start sites, generating truncated or out-of-frame neoproteins. We uncover a fusion-dependent increase in ribosomal readthrough into poly(A) tails, generating a stable 80-amino-acid TRPM4 neoprotein that, despite lacking a stop codon, is the most abundant tumor-specific microprotein across patients. Proteomic, immunopeptidomic, and immunofluorescence analyses validate neoproteins as tumor-restricted antigens, revealing a therapeutically actionable cancer dark proteome controlled by one oncogenic fusion.

**Highlights:**

- Fusion proteins activate in- and out-of-frame neoproteins through intragenic transcription.
- Neoproteins exhibit distinct protein structures and localizations compared to their host genes.
- Mass spectrometry detects 911 cryptic proteins, including 29 EwS neoproteins, 15 HLA-I presented.
- TRPM4-dORF is a stop-less 80-aa neoprotein with abundant peptide evidence exclusively in EwS.

## Introduction

Ewing sarcoma (EwS) exemplifies how chromosomal translocations encoding chimeric fusion oncoproteins with aberrant transcription factor (TF) activity can reshape entire transcriptomes. This pediatric bone and soft tissue sarcoma is driven by fusions between members of the *FET* gene family (*FUS*, *EWSR1*, *TAF15*) and *ETS* (E26 transformation-specific) family TF genes, most commonly *EWSR1::FLI1* (EF1; 85% of cases), which create chimeric TFs with neomorphic DNA-binding properties^1–5^. The resulting fusion proteins gain affinity for GGAA microsatellite repeats, sequences absent from the normal *ETS* binding repertoire^6,7^. Fusion protein binding at GGAA microsatellites opens chromatin and drives transcription of nearby genes, activating oncogenic programs through EwS master regulators like *NR0B1*, *SOX6* and *NKX2-2*^8–14^. However, GGAA microsatellites are not conserved across species’ genomes and their polymorphisms have been shown to contribute to susceptibility to EwS^15–18^.

Beyond activating known oncogenes, fusion binding occurs throughout constitutively silent genomic regions, including intergenic spaces and gene introns. This aberrant epigenetic change can transcriptionally activate previously unannotated sequences we termed neogenes^19^. When binding occurs close to or within canonical genes, newly induced transcription start sites (TSSs) or local enhancers can induce a multitude of alternative transcript isoforms (neoisoforms) that are similarly specific to EwS, as showcased by reports demonstrating pro-proliferative and/or pro-metastatic effects of *LOXHD1*^20,21^ and *KCNN1*^22^ neoisoforms.

Neogene translation signatures resemble those of translated non-canonical open reading frames^23–25^ (ncORFs, hereafter referred to as nc-translons^26^), which are defined as ribosome-decoded regions^26^: they are typically short, evolutionarily young and highly tissue/tumor-specific, explaining their absence from standard annotations. Crucially, nc-translons can encode microproteins with functional roles in physiology and disease, including cancer^27–32^. Nonetheless, we lack functional understanding of the putative biological roles of neogenes and the small proteins they produce, termed neoproteins. Notably, it remains unclear whether neoproteins impact cell biology and whether they evolved through similar mechanisms as other evolutionarily recent genes and nc-translons.

Neogenes and neoisoforms present unique therapeutic opportunities due to their tumor-specific expression. Their translation into neoproteins may generate HLA I-presented peptides as tumor-specific antigens. Because expression is directly driven by the oncogenic fusion, such targets could enable cancer vaccines with limited immune escape potential, addressing the urgent need for new treatments. However, several critical unknowns persist. As our prior studies^19^ were performed on relatively small numbers of in vitro models, neoprotein expression recurrence across EwS patients remains unknown. Furthermore, the relationship between neoprotein-encoding RNA abundance and HLA-I presentation of the protein product has also not been explored yet. This is essential information for pushing neoprotein therapeutics to the clinic.

To globally deconvolute complex neogene and neoisoform structures, map translated ORFs (translons) within these transcripts, and provide conclusive evidence for the produced neoproteins and other dark proteome constituents, we designed a framework that integrates long-read-informed transcriptome reconstruction, deep ribosome profiling (Ribo-seq), evolutionary genetics, and mass spectrometry-based proteomics and immunopeptidomics.

Across 48 patient tumor tissues, we identified and characterized 124 intergenic neogenes as well as 2,244 neoisoforms activated through aberrant fusion binding within 252 canonical genes. Many of these host small translated EwS-specific translons, which we assessed for recurrence across patients.

Compiling global proteomes matching tumors profiled with RNA-seq and Ribo-seq, a large set of EwS and non-EwS cell lines^19^, a deep multiprotease dataset of the A-673 cell line^33^, as well as EwS immunopeptidomes^34^, we detect endogenous peptide evidence for 911 nc-translons, including 13 neoproteins originating from neogenes and 16 originating from neoisoforms. Within this set, we find 15 neoproteins presented on HLA-I molecules: 11 from neogenes and 4 from neoisoforms. Overexpression and endogenous tagging of 19 putative neoproteins confirmed the stable expression and subcellular localization for 8 neoproteins, including an N-terminally truncated neoprotein produced from the usually meiosis-restricted *KASH5* gene, showcasing a new mechanism of cancer-testis antigen reactivation.

Our work establishes a comprehensive resource for understanding how fusion-driven pediatric cancers activate, and possibly functionally exploit, the cancer dark proteome, paving the way for clinical studies toward the safety and efficacy of neoprotein-targeting cellular therapies and therapeutic vaccines in children and young adults with EwS.

## Results

### Neogenes and neoisoforms are widespread and recurrently expressed across EwS patients

To map the full repertoire of EwS-specific genes and transcripts, we built a reference-guided transcriptome assembly combining short-read RNA-seq from 133 EwS patients with two independent long-read technologies (Oxford Nanopore direct RNA-seq and PacBio HiFi cDNA sequencing) from four EwS cell lines plus one EF1-fusion-engineered mesenchymal stem cell (MSC) line^35^ (**Figure 1a, S1a-e**). We then performed a series of rigorous filtering steps tailored to the known biology of EwS to identify EwS-specific neogenes and neoisoforms.

**Figure 1.**
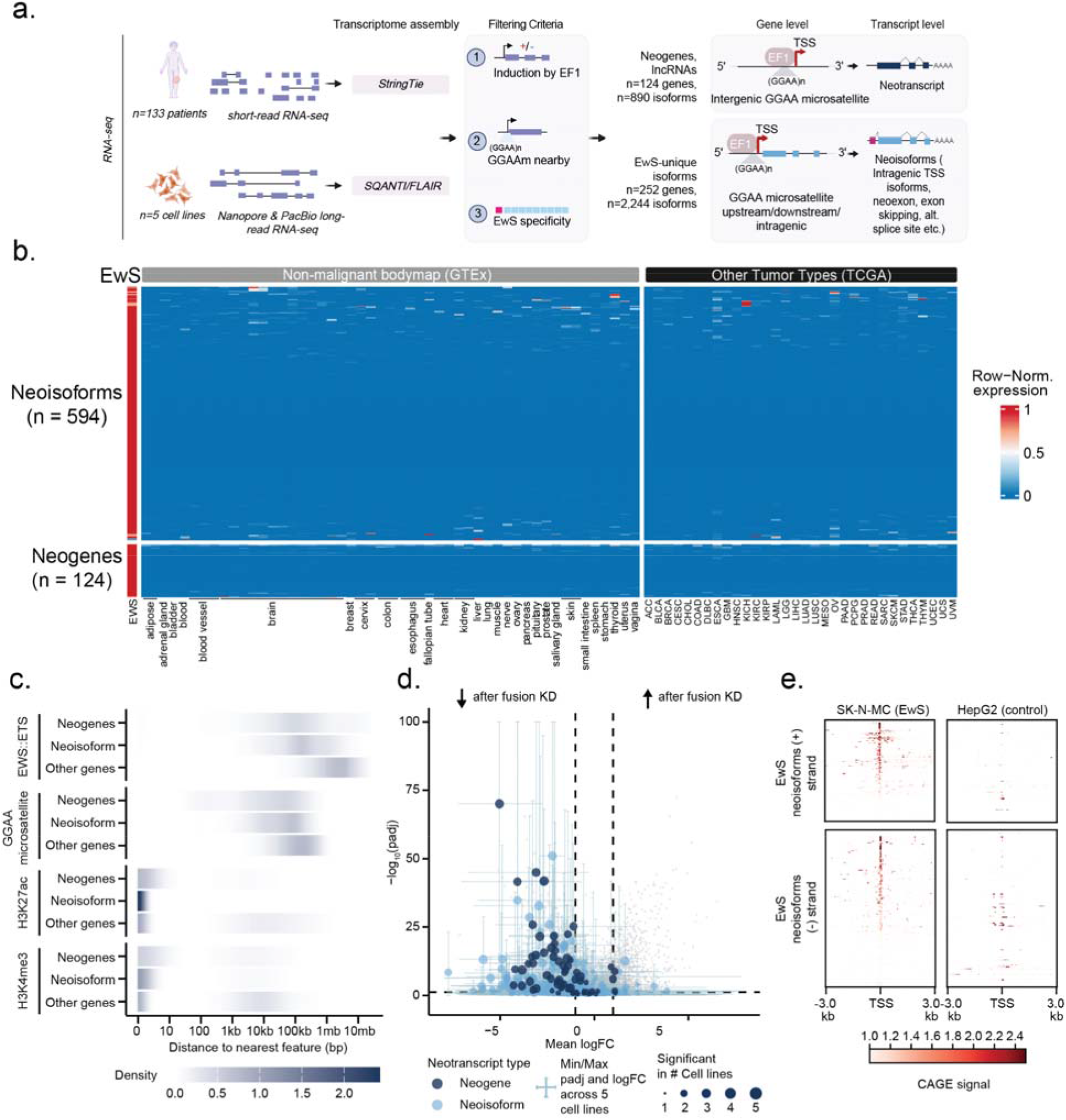
Integration of short- and long-read RNA-seq reveals EwS specific neogenes and neoisoforms. **a**. Graphical overview of the short- and long-read RNA-seq samples, pipelines used for processing (see Methods) an the different EwS-specific transcript types. **b**. Heatmap showing row-normalized expression values, with each row being a neogene or a neoisoform. For the neoisoforms, only the transcript sequences that are unique to EwS and unannotated were queried. See Methods for details. **c**. Distance distributions from TSSs to nearest *EWSR1::ETS* binding sites, GGAA microsatellites, H3K27ac peaks, and H3K4me3 peaks for neogenes, neoisoforms, and other genes. Neogenes and neoisoforms show significant enrichment at all regulatory features (Kolmogorov-Smirnov test, all adjusted p < 1 × 10^-74^). ChIP-seq data from Orth et al., 2022^10^. **d**. Volcanoplot exhibiting average change i expression after dox-induced shRNA-mediated fusion knockdown across 5 different EwS cell lines (4x *EWSR1::FLI1*, 1x *EWSR1::ERG*). Neogene and neoisoform are separated by color, the size of the dot indicates the number of cell lines in which a transcript was changed significantly. Light blue whiskers illustrate minimum and maximum values for p_adj_ and logFC values across 5 cell lines for each transcript. **e**. CAGE-data signal at the TSS of EwS neoisoforms in SK-N-MC EwS cell line and HepG2 control Hepatoblastoma cell line, highlighting EwS-specific TSS usage.

We selected genes downregulated upon EF1 knockdown (KD) in cellular models, in proximity to GGAA microsatellites, and specifically expressed in EwS cells and tumors (**Figure 1a**). We then cross-referenced our transcriptome against a comprehensive panel of publicly available RNA-seq datasets encompassing: 17,350 non-malignant tissue samples across 52 tissue types from the Genotype Tissue Expression project (GTEx); 12,665 EF1-negative malignant samples spanning 33 adult cancer types (TCGA) and 10 pediatric cancer types (St. Jude Children’s Research Hospital and Princess Máxima Center for Pediatric Oncology); developmentally diverse non-malignant tissues from the Evo-devo atlas^36^; and an MSC reference panel. This multi-cohort filtering strategy revealed 890 transcripts arising from 124 fully intergenic neogene loci and 2,244 neoisoforms embedded within 252 annotated protein-coding genes, all exhibiting EwS-restricted expression (**Figure 1a–b**; **Supplementary Table S1-S3**). Of note, because accurate quantification of neoisoforms is hindered by homologous sequences flanking EwS-specific sequences, expression could be reliably estimated for all 124 fully intergenic neogenes and only a subset of 594 (out of 2,244) neoisoforms with EwS-specific sequences. We termed such fully unique exonic sequences within neoisoforms “neoexons” throughout this study (**Figure 1b**, see Methods).

Analysis of published EWSR1::ETS, H3K27ac and H3K4me3 ChIP-seq data from the Ewing Sarcoma Cell Line Atlas (ESCLA)^10^, alongside GGAA microsatellite locations confirmed that neogene and neoisoform TSSs localize significantly closer to these regulatory features compared to other genes (Kolmogorov-Smirnov test, p_adj_ < 1 × 10^-74^; **Figure 1c**). To functionally confirm fusion-dependance, we performed RNA-seq on five cell line models with doxycycline (dox)-inducible *EWSR1::ETS* fusion knockdown, in triplicates. A-673 cells expressing a scrambled short hairpin RNA (shRNA) (A-673/TR/shControl) showed minimal transcriptional changes upon dox treatment, confirming specificity of the knockdown effect, whereas shRNAs targeting EWSR1::FLI1 drastically reduced the expression levels of neogenes and neoisoforms across all models tested including the TC-106/TR/shERG cell line harboring an *EWSR1::ERG* fusion **(Figure 1d, Figure S2a-b)**. This pattern distinguishes EwS-specific transcription from general cancer-associated transcripts and establishes their direct dependence on fusion protein activity.

To independently validate the neoisoform TSSs defined by long-read sequencing, we analysed Cap Analysis of Gene Expression (CAGE) data from the EwS cell line SK-N-MC (FANTOM^37,38^). Because CAGE maps capped 5′ transcript ends at base-pair resolution, it resolves precise promoters and TSSs of individual transcripts. CAGE signal accumulated at the predicted neoisoform TSSs in SK-N-MC, but was largely absent in the non-EwS hepatoblastoma HepG2 cell line (**Figure 1e**), confirming these as bona fide EwS-restricted start sites.

Combined, this strategy significantly expanded our previous neogene set^19^ from 26 to 124 neogenes and, importantly, uncovered a previously uncharacterized layer of transcriptomic-driven complexity with the identification of 2,244 novel EwS–specific neoisoforms that may contribute to the production of EwS-specific neoantigens.

##### Box 1. Nomenclature

**Translon** —

A ribosome-decoded genomic region, proposed as a unifying term for any translated sequence regardless of its coding potential or annotation status^26^. In this study, we use translon to refer to both canonical and non-canonical translated ORFs identified by ribosome profiling.

**Coding DNA sequence (CDS)** —

The portion of a gene’s mRNA that is translated into protein, spanning from the annotated start codon to the stop codon. In this study, CDS refers to the canonical, reference-annotated coding sequence of a protein-coding gene.

**Non-canonical translon (nc-translon)** —

A translon that does not correspond to an annotated protein-coding CDS. Nc-translons include upstream ORFs (uORFs), downstream ORFs (dORFs), translons in lncRNAs, and other previously unannotated translated regions.

**Neogene** —

A fusion-driven, EwS-specific gene arising from a normally silent (intergenic) genomic locus that is transcriptionally activated through EWSR1::ETS binding at GGAA microsatellites^19^.

**Neoisoform** —

An EwS-specific transcript isoform of a known protein-coding gene, generated through fusion-induced mechanisms such as neoexon inclusion, intragenic TSS activation, or alternative splicing. Neoisoforms can encode truncated in-frame or entirely out-of-frame protein products relative to the canonical CDS.

**Neoexon** —

An exonic sequence transcribed from a canonical protein-coding gene, absent from GENCODE transcriptomic reference and specifically expressed in EwS.

**Neoprotein** —

Any protein product encoded by a neogene or neoisoform translon. Neoproteins are tumor-restricted and include both truncated variants of known proteins (from in-frame neoisoforms) and entirely novel protein sequences (from neogenes and out-of-frame neoisoforms).

**Microprotein** —

A small protein (typically <100 amino acids) encoded by a non-canonical ORF, with sufficient evidence for stable expression and functional relevance to warrant annotation as a protein-coding gene.

**Peptidein** —

A translation product that can be demonstrated to exist but does not meet the criteria for protein annotation due to insufficient evidence for a physiological role. They lack convincing evolutionary or homology-based support for function and are often identified through immunopeptidomics or through proteomics data derived from cancer samples or cell lines^39^.

**Dark proteome** —

The collective set of proteins and peptides encoded outside of annotated protein-coding sequences, encompassing microproteins, peptideins, and other nc-translon-derived products. These are typically absent from standard reference proteomes.

### Recurrent cancer-specific translation across 48 EwS patients

Next, we sought to assess the translational potential of EwS-specific neogenes and neoisoforms by determining the fraction of their RNA molecules bound by ribosomes (ribosome-protected fragments, RPFs), indicative of neoprotein synthesis. We performed ribosome profiling across 48 fresh-frozen EwS patient tumor samples. In parallel, we profiled 5 patient-derived cell line models with and without a dox-inducible shRNA targeting the fusion protein, from our previously published ESCLA^10^, in triplicates **(Figure 2a, Supplementary Table S4)**. Our analysis yielded more than 4 billion RPFs, making it one of the deepest translatome resources to date and the single largest pediatric cancer translatome **(Figure S3a-b)**. Samples exhibited excellent three-nucleotide codon periodicity (average 84% for 29-nt RPFs) **(Figure 2b, Figure S3c)**, stressing the high predictive value for identifying new translons.

**Figure 2.**
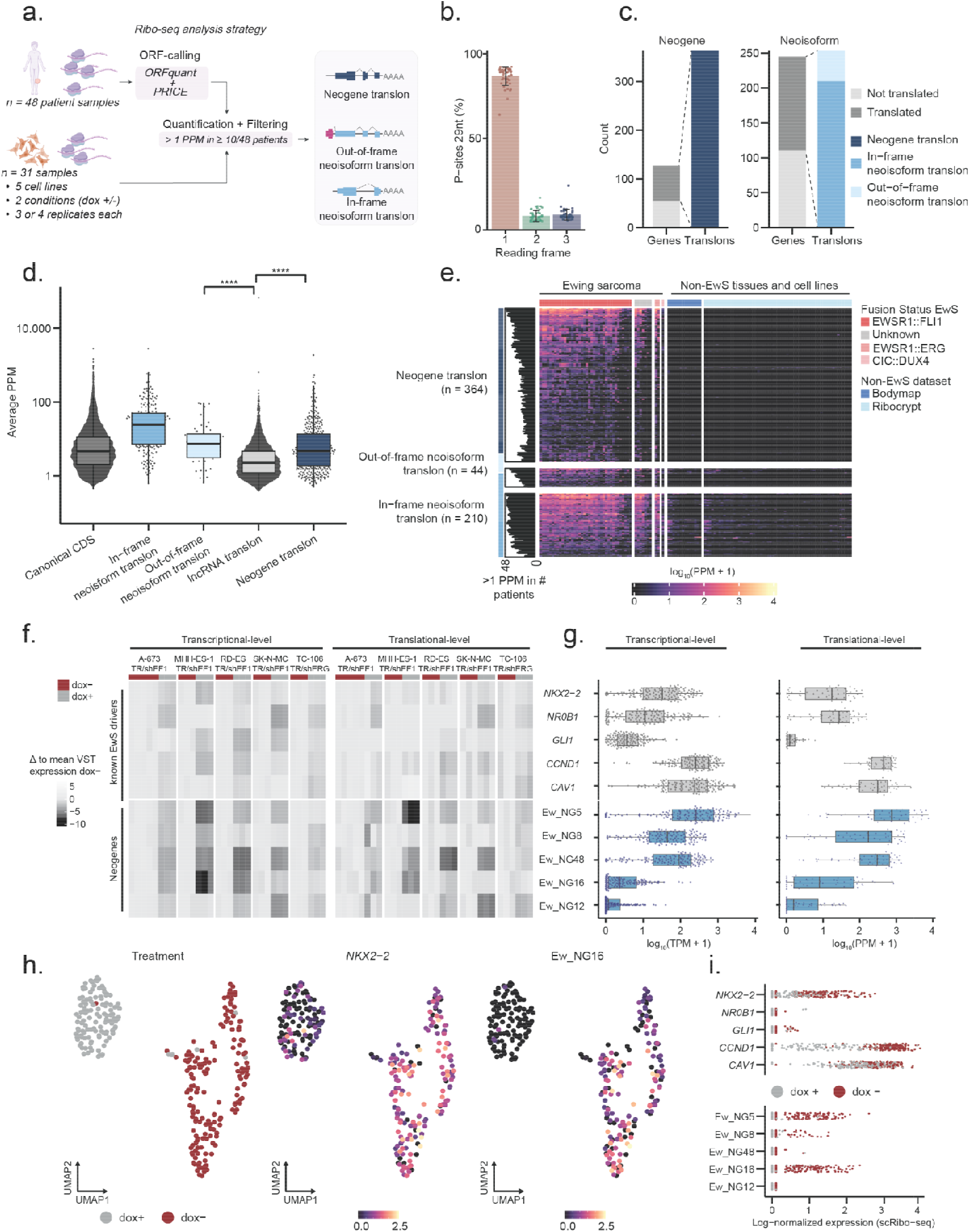
Deep bulk and single cell ribosome profiling reveals widespread and abundant translation of EwS-specific neogenes and neoisoforms. **a.** Overview of Ribo-seq processing and filtering strategy for 48 patients and 5 cell line samples to identify neogene, in-frame and out-of-frame neoisoforms translons. **b.** Percentage of patient sample ribosome-protected fragments with 29nt length attributed to each reading frame of canonical genes. **c.** Stacked barplot showing the number of translated transcripts with the numbers of translons. Color indicates the translon-type. **d.** Average in-frame P-sites per million (PPM) for canonical-, lncRNA-, neogene-, and neoisoform translon types. PPM plotted on a log10 scale. **** indicates p ≤ 0.0001. **e.** Heatmap depicting log (PPM + 1) for each neogene or neoisoform translon across patient samples, split by fusion type and in non-EwS Ribo-seq datasets from the Bodymap^23^ and RiboCrypt. The barplot on the left indicates the number of patients with >1 PPM for each translon. **f.** Heatmap showing the change in expression relative to mean dox-negative values after dox-induced fusion knockdown in 5 EwS cell lines, for 5 well-established fusion targets and 5 neogenes at the transcriptional (RNA-seq, left) and translational (Ribo-seq, right) level. **g.** Transcriptional (log (TPM + 1)) and translational (log (PPM + 1)) levels of 5 well-established fusion targets and 5 neogenes across patient samples. **h.** UMAP projections of single-cell ribosome profiling (scRibo-seq) data from MHH-ES-1/TR/shEF1 cells, colored by dox treatment, *NKX2-2* expression, and Ew_NG16 expression. **i.** Log-normalized scRibo-seq expression values for 5 well-established fusion targets and 5 neogenes, split by dox treatment.

We used two complementary translon-calling approaches (ORFquant^40^ and PRICE^41^), followed by quantification of a unified translon list across all 48 patients, employing stringent filtering for confirmed in-frame ribosome occupancy in at least 10 of 48 patients with a P-sites per million (PPM) cutoff of greater than 1. This yielded 364 neogene translons mapping to 72 out of 124 intergenic neogenes (∼58%), and 254 neoisoform translons mapping to 121 out of 252 protein-coding genes with detected neoisoforms (∼53%) **(Figure 2c, Supplementary Table S5)**.

Translation of neoisoforms in the 121 annotated coding genes was categorizable into two types of translons. First, for 210 neoisoform translons in 111 protein coding genes, we identified in-frame EwS-specific alterations, each producing EwS-unique coding sequence variations of the annotated CDS. Second, 23 genes generated EwS-specific neoisoforms encoding 44 out-of-frame translons, producing protein sequences entirely different from any canonical CDS and not previously described in EwS or any other malignancy **(Figure 2c)**. These two separate neoisoform translon classes highlight how fusion-driven transcriptomic changes can lead to distinct translatomic outcomes previously largely unknown in EwS.

The depth of our dataset further enabled the detection of 24,478 non-canonical translons beyond those classifying as EwS-restricted, spanning uORFs, lncRNA translons and translons in additional unannotated genomic regions not specific to EwS **(Figure S3d)**. Of these, 11,069 (44.6%) had in-frame overlap with translons from recent large-scale cataloging efforts^42^, not only demonstrating the validity of our data but also its potential for extending future reference annotations **(Figure S3e)**.

### Neogene translation far exceeds lncRNA translation in EwS patients

Despite being transcribed and translated from normally non-coding DNA, EwS-specific translons are translated at strikingly high levels. Neogene translons and out-of-frame neoisoform translons showed median translation rates broadly comparable to canonical genes and exceeded those of translated lncRNAs **(Figure 2d)**, while in-frame neoisoform translons reached even higher median levels. This is consistent with their partial retention of established coding sequence context, often overruling the translation rates of the local full-length CDS. For all neogene and neoisoform classes combined, translation patterns were consistent between *EWSR1*::*FLI1* and *EWSR1::ERG* fusion subtypes but largely absent in a patient with *CIC*::*DUX4* sarcoma, a morphological mimetic of EwS also belonging to the entity group of undifferentiated small round cell sarcomas of soft-tissue and bone^43^. This further highlights that EwS and *CIC::DUX4* sarcomas are distinct entities^44^ **(Figure 2e)**. As expected, neogene and neoisoform translons were completely absent from the RiboCrypt and Human Body Map resources^23^, covering almost four thousand Ribo-seq datasets of non-EwS tissues and cell lines, again confirming their specificity to the fusion-driven tumor.

The top-expressed neogenes (Ew_NG5, 8, 12, 16 and 48) showed marked transcriptional and translational downregulation following dox-induced shRNA-mediated fusion knockdown, exceeding the degree of downregulation observed for five well-characterized fusion targets (*CAV1, CCND1, GLI1, NKX2-2,* and *NR0B1*)^8,9,45–47^ **(Figure 2f)**. Moreover, in patient samples, mRNA abundance and translation rates of these five neogenes equaled and sometimes exceeded those of the well-established fusion targets **(Figure 2g)**. With translation levels reaching over 500 PPM, this, to our knowledge, significantly exceeds translation rates of any nc-translon in any lncRNA observed to date.

To evaluate whether neogene translation shows heterogeneity at the single-cell level, we performed single-cell ribosome profiling scRibo-seq^48^ on MHH-ES-1/TR/EF1 cells. Translation patterns on the single cell level correlate well with the bulk data (Spearman ρ = 0.50), validating our experimental setup **(Figure S4a-e)**. Because of the inherent sparsity of single-cell data, we merged the 364 neogene translons back into 72 neogenes for quantification, of which 31 were detected in MHH-ES-1/TR/EF1**(Figure S4f).** Top-expressed neogene translons displayed clear separation between dox-treated conditions **(Figure 2f-g, Figure S4g)**, and per-cell expression mirrored that of the *NKX2-2* CDS, with translation greatly reduced upon fusion knockdown **(Figure 2f)**, confirming fusion-dependency and generally homogeneous cancer cell expression at single-cell resolution.

Together, these findings establish widespread, recurrent, and abundant translation of fusion-dependent transcripts across patient tissue bulk, and single-cell EwS translatomes, with top-neogenes reaching translation levels in patients comparable to or exceeding established fusion targets.

### Neoisoform transcription from intragenic TSS and neoexon inclusion drives EwS-specific in- and out-of-frame translation

To elucidate the origins and mechanisms of neoisoform transcription and translation in EwS, we performed an in-depth analysis of neoisoform structures and features with long-read RNA sequencing. We classified the 2,244 EwS-specific neoisoforms observed in 252 protein coding genes, of which 550 neoisoforms in 121 genes contain translons, according to three distinct mechanisms (**Figure S5a-b, Supplementary Table S6**): (i) neoexon inclusion (ii) intragenic TSS induction and (iii) other alternative splicing isoforms.

First, we identified 594 neoisoforms across 111 genes carrying neoexons, which we define as novel exonic sequences absent from any annotated transcript. In total, 149 neoisoforms in 51 genes contain translons detected by Ribo-seq, of which 76 result in truncated but in-frame neoprotein production and 15 produce out-of-frame neoproteins **(Figure S5c)**. Two examples illustrate this neoexon mechanism driving neoprotein formation **(Figure 3a-b)**: At the *TNS1* gene locus, a novel intragenic TSS, also corroborated by SK-N-MC CAGE-data, generates an EwS-unique starting neoexon producing two out-of-frame neoisoform translons with clear translation signals detectable in 31/48 and 34/48 patients **(Figure 3a)**. In the well-established EwS fusion target gene *NKX2-2*^9^, an EwS-specific neoisoform encodes an out-of-frame translon located upstream of the annotated exon 1, showing recurrent ribosome occupancy across 42/48 patients **(Figure 3b)**.

**Figure 3.**
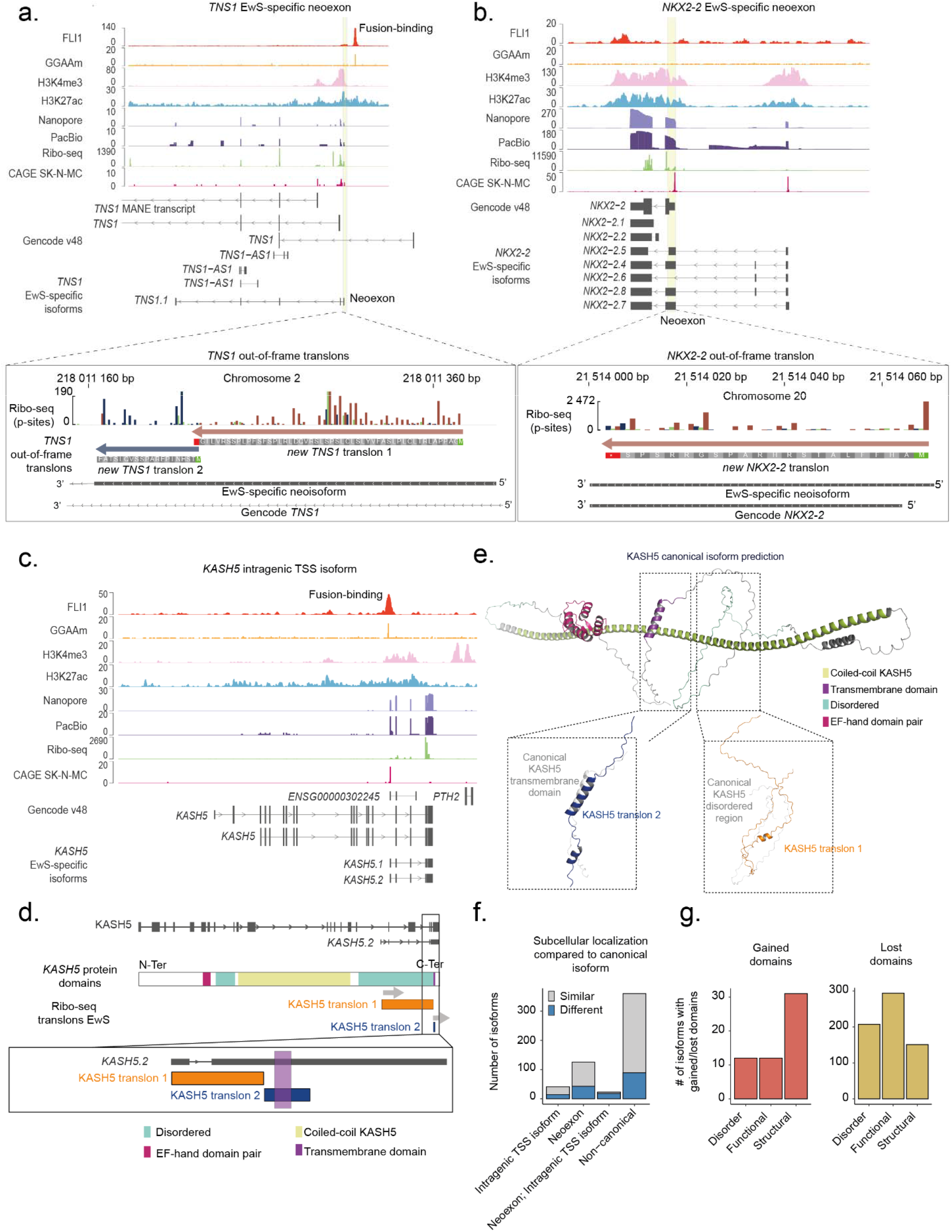
Long-read RNA sequencing reveals distinct mechanisms driving in- and out-of-frame neoisoform translation. **a.** Genomic visualization of the *TNS1* gene, depicting its EwS-specific isoform. The yellow highlighted genomic sequence represents a novel transcribed sequence that encodes two out-of-frame neoisoforms translons. A zoom in this neoexon shows the two out-of-frame translons Ribo-seq quantification where the P-sites are colored by frame. **b.** Genomic overview of the *NKX2-2* gene, showing its canonical and EwS-specific neoisoforms. In EwS, neoisoforms transcription starts upstream of the canonical *NKX2-2* first exon and encode for an out-of-frame EwS-specific translon. The yellow-highlighted region shows the out-of-frame translon being translated from the neoisoform only. P-sites are colored by frame. **c.** Genomic view of two *KASH5* intragenic TSS isoforms associated with EF1 fusion binding at GGAA microsatellites. Nanopore, PacBio, and Ribo-seq coverage is restricted to the last exons of the *KASH5* gene. **d**-**e**. Schematic of (d) protein functional domains and (e) alphafold predictions for KASH5-derived neoproteins. The intragenic TSS-derived *KASH5.2* isoform gives rise to two Ribo-seq translons highlighted in two zoom-in regions, where *KASH5* translon 2 (MSC-EF1_00035195_394_555) retains the KASH5 transmembrane domain, whereas *KASH5* translon 1 (MSC-EF1_00035195_34_390) retains only disordered regions. **f**. Barplot showing the number of neoisoforms with at least one translon with a different DeepLoc predicted subcellular localization in comparison with its canonical counterpart. **g**. Barplot showing the number of neoisoforms (all categories included) giving rise to at least one translon with a predicted gained or lost domain in comparison with its canonical isoform counterpart.

Second, we identified 321 neoisoforms across 32 genes expressed through a *de novo* intragenic TSS, induced by *EWSR1::ETS* binding at nearby GGAA microsatellite repeats. Of note, 155 out of 321 intragenic TSS isoforms lead to the transcription of a fully unique neoexon, creating redundancy with the first category described above **(Figure S5a-b)**. A striking example is the aberrant activation of the 3′ part of the meiosis-restricted *KASH5* gene, representing a new mechanism of cancer-testis antigen (CTA) activation. In *KASH5*, long-read RNA-seq and Ribo-seq coverage was restricted to the aberrantly active 3′ region of the gene, located downstream of an intragenic EF1-bound GGAA microsatellite. CAGE-data also confirms this novel TSS to be highly active in EwS. EF1 binding induced two neoisoforms (**Figure 3c-d)**, of which the *KASH5.2* isoform gives rise to a 54-aa (translon 2) and a 119-aa translon (translon 1). Both translons lose all coiled-coiled and (EF-hand) helix-loop-helix domains typical for the canonical KASH5 structure, as shown with AlphaFold3 structure prediction (**Figure 3e)**. The C-terminal transmembrane domain of KASH5 is retained in only one translon. However, due to N-terminal truncation, it lacks a signal peptide and is predicted to be extracellular with 0.997 likelihood by DeepLoc^49^, suggesting biomarker potential. The other translon is predicted to be fully disordered and has a 0.45 likelihood of cytoplasmic intracellular localization according to DeepLoc. At a mean PPM of 36.8, *KASH5* is among the most highly expressed CTA in EwS across 239 referenced CTAs in the CTDatabase and CTexploreR databases (**Figure S5d**)^50^.

The third and last category includes 1,484 EwS-specific neoisoforms that did not belong to either of the two previous categories and resulted from multiple alternative splicing events **(Figure S5e)**. These neoisoforms encoded for 158 in-frame and 31 out-of-frame translons **(Figure S5f, Materials and Methods)**.

Truncated neoprotein variants of known proteins are likely to lose one or more functional domains and, when N-terminally altered, are prone to change their subcellular localization. Indeed, out of 190 predicted neoproteins translated from neoisoforms generated through either neoexon formation or an intragenic TSS, 76 (40%) showed a different DeepLoc prediction from the canonical protein **(Figure 3f)**. In particular, 20 neoproteins (12 in-frame, 8 out-of-frame) across 6 genes were predicted to acquire cell membrane or extracellular localization unlike their canonical protein counterparts identifying them as particularly compelling candidates for extracellular therapeutic targeting. Similarly, using InterProScan^51^ domain prediction, we found that 361 neoisoforms across 80 genes lead to at least one translon with a loss of protein domain compared to its canonical counterpart while 43 neoisoforms in 8 genes exhibited an additional protein domain **(Figure 3g)**.

For each of the three categories of neoisoform translons, translation as measured by Ribo-seq was markedly reduced across 5 cell line models following dox-inducible shRNA-mediated knockdown of *EWSR1::ETS* **(Figure S5g)**. Notably, out-of-frame neoisoform translons showed significantly greater reduction compared to the full-length CDS of canonical isoforms from the same genes **(Figure S5h).** This indicates fusion control of the neoisoform translon while the canonical isoform was not affected, confirming the strong EwS specificity of neoisoform translons. Collectively, our data further expand the spectrum of aberrant transcriptional and translational consequences of *EWSR1::ETS* fusion oncoproteins.

### Detection of EwS-specific neoproteins in deep proteome and immunopeptidome data

As neogene and neoisoform translons represent a novel and attractive class of potential therapeutic targets, we investigated their detectability and abundance by mass spectrometry, either as cellular neoproteins or as peptides presented by HLA-I complexes on the cell surface. We collected four independent proteomics datasets^19,33,34^ spanning distinct sample types, digestion strategies, and acquisition methods, covering 122 samples including 86 distinct (tumor) tissues **(Figure 4a, Supplementary Table S7)**. We searched these spectra against a custom database derived from our combined analysis of short-, long-read RNA-seq, and ribosome profiling analyses containing neogene translons, in-frame and out-of-frame neoisoform translons, and 24,478 additional general nc-translons, with translon-type FDR control **(Figure S3e)**. In total, we found peptide evidence for 911 nc-translons, irrespective of whether these were restricted to EwS. Of these, 765 were found in immunopeptidomics data and 155 were found in at least one global proteomics dataset, with only 9 being detected in both EwS global proteomics and immunopeptidomics data. Of the 155 global proteome detections, 79 nc-translon protein products were found in more than one biological sample. Thirty-three of these were recurrently identified between independent global proteome datasets, and 55 had evidence from more than one unique peptide, potentially providing sufficient evidence for nominating these as putative new microproteins or peptideins^39^ (**Figure 4b, Supplementary table 7-8)**.

**Figure 4.**
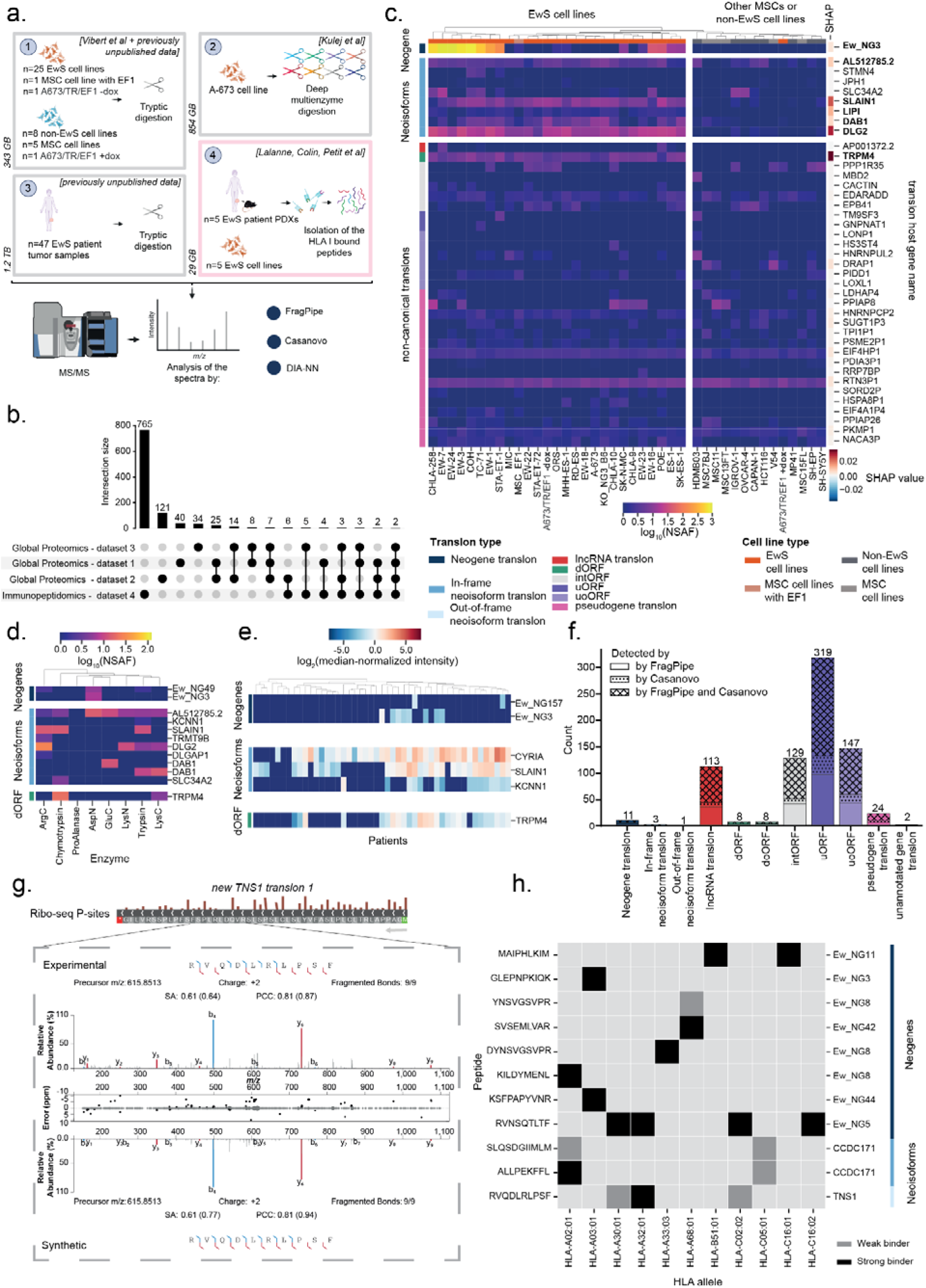
Proteomics detection of nc-translons and neoproteins across multiple datasets. **a.** Overview of proteomics datasets searched against the nc-translon and neoprotein database, including sample composition and data volumes. **b.** Upset plot showing inclusive intersections of nc-translon detections across all 4 datasets. Each bar represents the number of translons detected in that combination of datasets; translons detected in multiple datasets contribute to multiple bars. Total unique detections: 911. **c.** Heatmap of nc-translons detected across 25 EwS cell lines and controls in dataset 1^19^ and previously unpublished data. Values represent log-transformed Normalized Spectral Abundance Factors (NSAF). Cell lines and translons were grouped by hierarchical clustering. Translon types and cell line identities are indicated by color-coded annotations (left). SHAP-values are displayed as color-coded annotations (right). See **Figure S6a** for details. **d.** Neogenes and neoisoforms detected in the A-673 deep multienzyme digestion dataset 2^33^. Values represent log-transformed NSAF. **e.** Neogenes and neoisoforms detected across 47 EwS patient tumor samples from previously unpublished dataset 3. Values represent log-transformed median-normalized intensity. **f.** Numbers of nc-translons detected per translon class in the immunopeptidomics dataset 4^34^. Hatching indicates translons confirmed by both Fragpipe database search and Casanovo *de novo* sequencing. **g.** Proteomics validation of a novel out-of-frame neoisoform of TNS1. Ribo-seq P-sites and predicted translons are shown alongside the MS/MS spectrum of the detected peptide. The endogenous peptide spectrum is shown on top, while the synthetic peptide spectrum is shown on the bottom. **h.** Peptides from neogenes and neoisoforms detected in dataset 4 and their HLA-restriction.

Searching global proteome LC-MS spectra of 27 EwS and 14 non-EwS cell lines^19^, we identified peptides supporting 1 neogene translon, 8 in-frame neoisoform translons, and 31 nc-translons, most of which are not specific to EwS **(Figure 4c)**. Detection of neoproteins from neogene Ew_NG3 and 5 out of 8 neoisoform translons was strictly specific to EwS samples. Despite the limited size of this detected set, hierarchical clustering cleanly separated EwS from non-EwS samples, and SHAP-based (Shapley Additive Explanations) feature importance analysis of a Random Forest classifier confirmed that neoisoform translons dominated the top-ranking features driving this classification **(Figure 4c, Figure S6a)**. To increase proteome coverage, we next searched a deep multienzyme digestion dataset of the EwS A-673 cell line^33^ **(Figure 4d, Figure S6b)**, detecting peptides matching 2 neogene translons, 9 in-frame neoisoform translons, and 111 nc-translons not specific to EwS.

To complement both cell line profiling studies, we additionally generated a new global proteomics dataset from a cohort of 47 EwS patient tumor samples, matching samples from 47 out of 48 patients used for Ribo-seq **(Figure 4e, Figure S6c)**. In these data, we detected proteins from 2 neogene translons, 3 neoisoform translons, and 29 nc-translons not specific to EwS, confirming that neogene-derived peptides can be directly observed in patient tumors.

Across all global proteome datasets, several neoisoform translons were reproducibly assigned as lead proteins by the parsimonious inference algorithm based on the Occam’s razor principle^52^, including truncated isoforms of SLAIN1 and KCNN1. As their supporting peptides are shared with canonical UniProt entries, these detections are consistent with elevated expression of the C-terminal portion of these proteins in EwS, corroborated by the Ribo-seq evidence confirming neoisoform translation across patient umors.

### Neogenes and neoisoforms are a rich source of tumor-restricted immunopeptides

Next, we interrogated HLA-I immunopeptidomics data from patient-derived cell lines and xenografts (PDX)^34^, as shown in **Figure 4f**. Despite being the smallest dataset in terms of total spectral matches, HLA-I presented peptides contributed most nc-translon-derived peptide evidence (765 out of 911 total detected nc-translons, of which 756 nc-translons were uniquely detected in immunopeptidomics data).

The HLA-I immunopeptidomics data yielded more neoprotein and neoisoform derived peptides than any of the global proteomics datasets, even though it was considerably smaller. We detected 11 neogene and 4 neoisoform proteins on HLA-I, versus only 1–3 neogene and 3–9 neoisoform proteins in each global proteomics dataset. This enrichment is consistent with the rapid processing and presentation of neoproteins hypothesis, as well as with recent reports^53,54^. This enrichment might partly be explained by the inherently low number of tryptic peptides produced by short neogene translons **(Figure S6d)** (median 5 vs 60 for canonical proteins), which limits their detection in standard proteomics. Accordingly, the three neogene translons detected in tryptic datasets ranked among the highest in predicted tryptic peptide count. In contrast, with a median of 15 predicted HLA strong binders per neogene translon, immunopeptidomics provides sufficient epitope diversity for reliable detection, whereas the median of 5 tryptic peptides makes global proteomics detection highly stochastic.

One of the most promising EwS-restricted and HLA-I presented tumor-specific neoantigens mapped to an out-of-frame neoisoform translon within the *TNS1* gene **(Figure 3a**, **Figure 4g)**. Manual curation of the Ribo-seq data confirmed the out-of-frame translation, and we could validate the peptide-spectrum match with a synthetic peptide **(Figure 4g)**. To further substantiate these results, we selected 2 additional peptide-spectrum matches mapping to 2 neoproteins for validation with synthetic peptides **(Figure S7)**. Discovered peptides and their HLA-restriction are shown on **(Figure 4h).**

In silico extrapolation of these high validation rates, population-level HLA allele frequencies, and the fact that each of the 5 EwS cell lines and 5 PDX models presented at least one nc-translon-encoded peptide on HLA-I, suggests that a neoprotein-based vaccine formulation would provide broad predicted coverage across diverse HLA backgrounds **(Figure S6e)**. This supports the potential utility of HLA-I presented neoproteins as a source of tumor-specific immunotherapy targets.

### The TRPM4-dORF is a stable neoprotein highly restricted to EwS that reveals potential EF1-mediated poly(A) translation readthrough

To our surprise, we were able to detect unique and EwS-restricted proteomics evidence for a non-canonical translon that we did not initially classify as EwS-specific, because it fell outside of our newly constructed neogenes and neoisoforms. This highly translated downstream translon (dORF) mapped to the 3′-UTR of the *TRPM4* gene, close to an EF1 bound GGAA microsatellite repeat **(Figure 5a-b)**. Up to five unique peptides mapping to this 80-codon long *TRPM4*-dORF were recurrently detected and present across global proteomics datasets, including in tryptic MS data of 27 out of 47 patient tumors, yet were completely absent from the non-EwS models **(Figure 4c-e**, **Figure 5a, Figure S6b)**. In fact, the *TRPM4*-dORF is the highest-ranking translon by SHAP analysis and thus a main determinant of the EwS classification within this dataset (**Figure 4c**).

**Figure 5.**
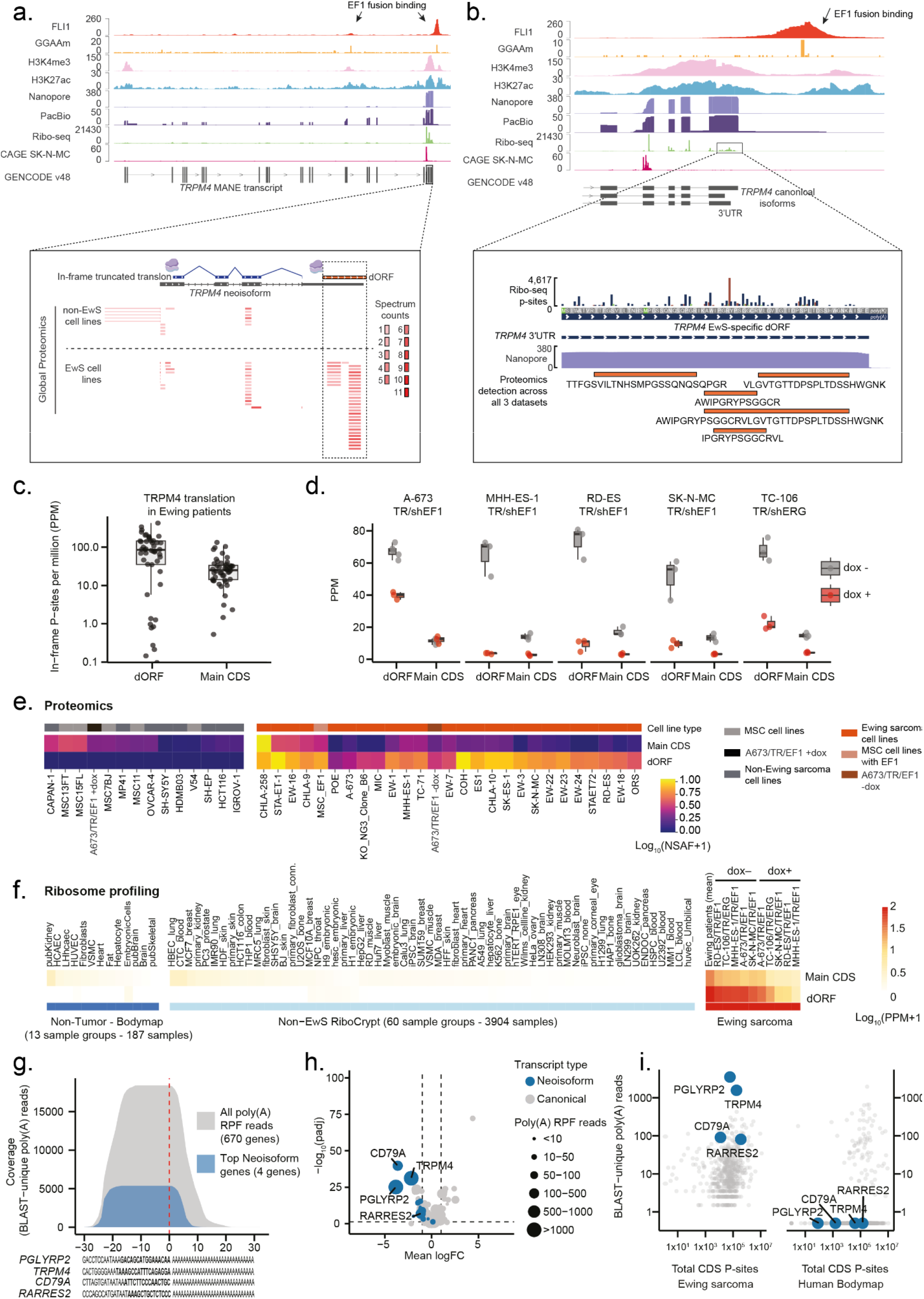
*TRPM4* harbors an EwS-specific downstream translon (dORF) driven by *EWSR1::ETS* activity that exemplifies poly(A) translational readthrough. a. Genomic overview of the full *TRPM4* gene depicting two FLI1-bound GGAA microsatellites and elevated long-read RNA-seq and ribosome coverage across the four terminal exons. Three *TRPM4* peptides mapping to *TRPM4*-dORF and identified by global proteome mass spectrometry from dataset 1 are annotated below the genomic track; each row represents an individual sample, with spectral counts represented as a color-coded vector. **b.** Genomic view of the *TRPM4* locus illustrating ribosome occupancy across the 3′ UTR of canonical isoforms, proximal to a FLI1-bound GGAA microsatellite. The highlighted region denotes an EwS-specific translon (dORF; TCONS_00433586_5495_5734) arising from 3′ UTR translation of *TRPM4* canonical transcripts. Ribo-seq P-sites are color-coded by reading frame, with the *TRPM4* EwS-specific nc-translon translated in the blue frame. No stop codon was found, instead the translation goes into the poly(A) tail. Orange boxes indicate the 5 peptides derived from the *TRPM4*-dORF detected in all 3 proteomics datasets^19,33,34^. **c.** Boxplots comparing translational output (Ribo-seq P-sites per million; PPM) of the *TRPM4* main CDS and dORF across EwS and non-EwS samples. **d.** Boxplots depicting translational levels (PPM) of the *TRPM4* main CDS and dORF in doxycycline-inducible EwS cell lines, in the presence or absence of dox-treatment. **e.** Heatmap summarizing quantification of the *TRPM4* main CDS and dORF by global proteomics across a mixed panel of EwS and non-EwS cell lines. **f.** Heatmap displaying translational levels of the *TRPM4* main CDS and dORF across a panel of non-tumor Ribo-seq datasets from the Human Bodymap and RiboCrypt compared to EwS Ribo-seq data. **g.** Coverage plot showing the number of BLAST-unique poly(A) junction RPF reads, with the coverage of the top neoisoform genes (*PGLYRP2*, *TRPM4*, *CD79A*, *RARRES2*) in blue. Below this panel, the sequence of each of the final 30nt is shown, with the minimal unique sequence in bold. **h.** Volcanoplot exhibiting average change in expression after dox-induced shRNA-mediated fusion knockdown across 5 different EwS cell lines (4x *EWSR1::FLI1*, 1x *EWSR1::ERG*). Genes with neoisoforms are highlighted in blue, the size of the dot indicates the number of poly(A) RPF reads found for each gene. **i.** Dotplot showing the number of BLAST-unique poly(A) reads vs their expression in either the EwS patients or in the non-malignant Human Bodymap dataset.

Notably, *TRPM4* showed a massive increase in RNA expression and ribosome occupancy in its final 4 exons as compared to the remainder of the gene body (**Figure 5b, Figure S8a**). Long-read RNA-seq data analyses indicated transcription from an alternative TSS giving rise to an unannotated isoform spanning the 4 last exons, with CAGE-data from SK-N-MC confirming this. Although this intragenic TSS isoform did not meet our inclusion criteria for EwS-specific neoisoform detection, it may explain the extremely high translation rates observed: in EwS patients, the *TRPM4*-dORF was translated ∼1.5-fold higher than the main CDS **(Figure 5c)**. It was also much more strongly downregulated than the main CDS upon *EWSR1::ETS* knockdown in five EwS cell lines with dox-inducible shRNAs against the respective fusions **(Figure 5d, Figure S8b)**, indicative of independent expression regulation. Strikingly, in a comparison with 60 different sample groups covering almost four thousand Ribo-seq datasets of the RiboCrypt and Human Body Map resources^23^, *TRPM4*-dORF translation was restricted to EwS despite the main CDS being translated ubiquitously (i.e., across non-EwS samples) on both the Ribo-seq and proteomics level **(Figure 5e-f)**. Notably, we observed background level translation rates (0.1 - 1 PPM) in pluripotent cell types, including H1, H9 and other embryonic stem cells, although expression levels in EwS were ∼100-fold higher.

When manually examining the *TRPM4*-dORF locus, we noted that the predicted dORF translation termination site falls outside of the transcript models supported by short- and long-read RNA-seq, as well as any GENCODE-annotated transcript isoform, suggesting translation into the poly(A) tail as part of likely ineffective non-stop decay **(Figure 5b)**. To confirm this, we searched RPF reads with a minimum of 5 trailing A’s mapping to the end of this transcript sequence uniquely (as determined by BLAST) and uncovered over a thousand poly(A) mapping RPF reads across our patient cohort (±1500) **(Figure 5g)**. To assess whether poly(A) readthrough translation extends beyond *TRPM4* as a broader phenomenon in cells and in EwS specifically, we mapped all Ribo-seq RPF reads from our EwS patient cohort and the Human Body Map resource to the last 30 nt 3′ UTR of all protein-coding transcripts, appended by 30 A’s. After filtering for uniquely mapped RPF reads, we found that in EwS two genes dominate the list of poly(A) readthrough reads, *PGLYRP2* and *TRPM4* **(Figure 5g-h, Figure S8c)**. Strikingly, genes with neoisoforms were well represented in the genes that showed abundant poly(A) readthrough translation. Despite the Human Bodymap^23^ containing more than double the number of RPF reads, it contained less total poly(A) readthrough RPF reads (15,779 vs 19,474 in EwS) within fewer genes (83 vs 670 in EwS). These results suggest that EF1 may influence poly(A) translation readthrough, especially for *PGLYRP2* and *TRPM4* genes for which poly(A) RPF reads are completely absent from any non-malignant tissue despite similar expression patterns **(Figure 5i)**.

Collectively, our proteomics analyses demonstrate that EwS-specific neoproteins are detectable across diverse platforms and clinical material, either as stable proteins in global proteomics or as HLA-I presented peptides in immunopeptidomics. The *TRPM4*-dORF exemplifies an unexpected mechanism to cancer-specific neoprotein production: it is translated from the 3′ UTR of a canonical transcript, likely through poly(A) readthrough of an otherwise non-stop transcript. That this phenomenon appears enriched among neoisoform-bearing genes and less present in non-malignant tissues raises the possibility that EF1-driven transcriptional rewiring not only generates new transcripts but also alters translational fidelity at existing loci, a mechanism that warrants further investigation. Together with the neogene and neoisoform translons characterized above, these findings reveal multiple, mechanistically distinct routes through which a single fusion oncoprotein expands the EwS-specific dark proteome.

### Most, but not all neoproteins are novel entities that emerged from neutrally evolving genomic sequences

Having established that neoprotein-derived peptides are detectable in global proteomics data and enriched on HLA-I, we next sought to characterize the full-size neoproteins they represent. Unlike in-frame neoproteins, which retain partial canonical protein architecture **(Figure 3)**, neoproteins encoded by neogene and out-of-frame neoisoform translons are expected to represent entirely novel protein sequences without any canonical counterpart. However, to understand their molecular properties and evaluate whether their evolutionary origins might reveal ancient gene reactivation, we combined protein domain prediction, sequence feature analysis, and phylogenetic reconstruction across 120 mammalian species.

To investigate when these neoprotein ORF sequences emerged and whether they represent remnants of ancient genes or recently arisen genomic sequences, we performed phylogenetic analysis using syntenic alignments across 120 species **(Figure S10a)**. Amino acid sequence similarity was high among primates but declined sharply in more distant species, indicative of a recent origin for most, in line with our earlier observations^24^ **(Figure 6a, Supplementary Table S9)**. Phylogenetic classification of intact orthologs across these species revealed that the majority of neogene (219 out of 364) and out-of-frame neoisoform translons (14 out of 44) arose within the primatomorpha clade, or more recently as exemplified by Ew_NG8 **(Figure 6b, Figure S9b-c)**, though a small number of neogene translons (9 translons from 4 neogenes) originated through pseudogenization **(Figure S9d)**. Length- and position-matched non-translated control ORF sequences (NT controls) showed similar sequence features and evolutionary origins **(Figure 6a)**, indicating that neogene and neoisoform translons at large reflect neutrally evolving genomic regions that stochastically acquired translation-competent ORFs, only to be aberrantly activated by fusion binding.

**Figure 6.**
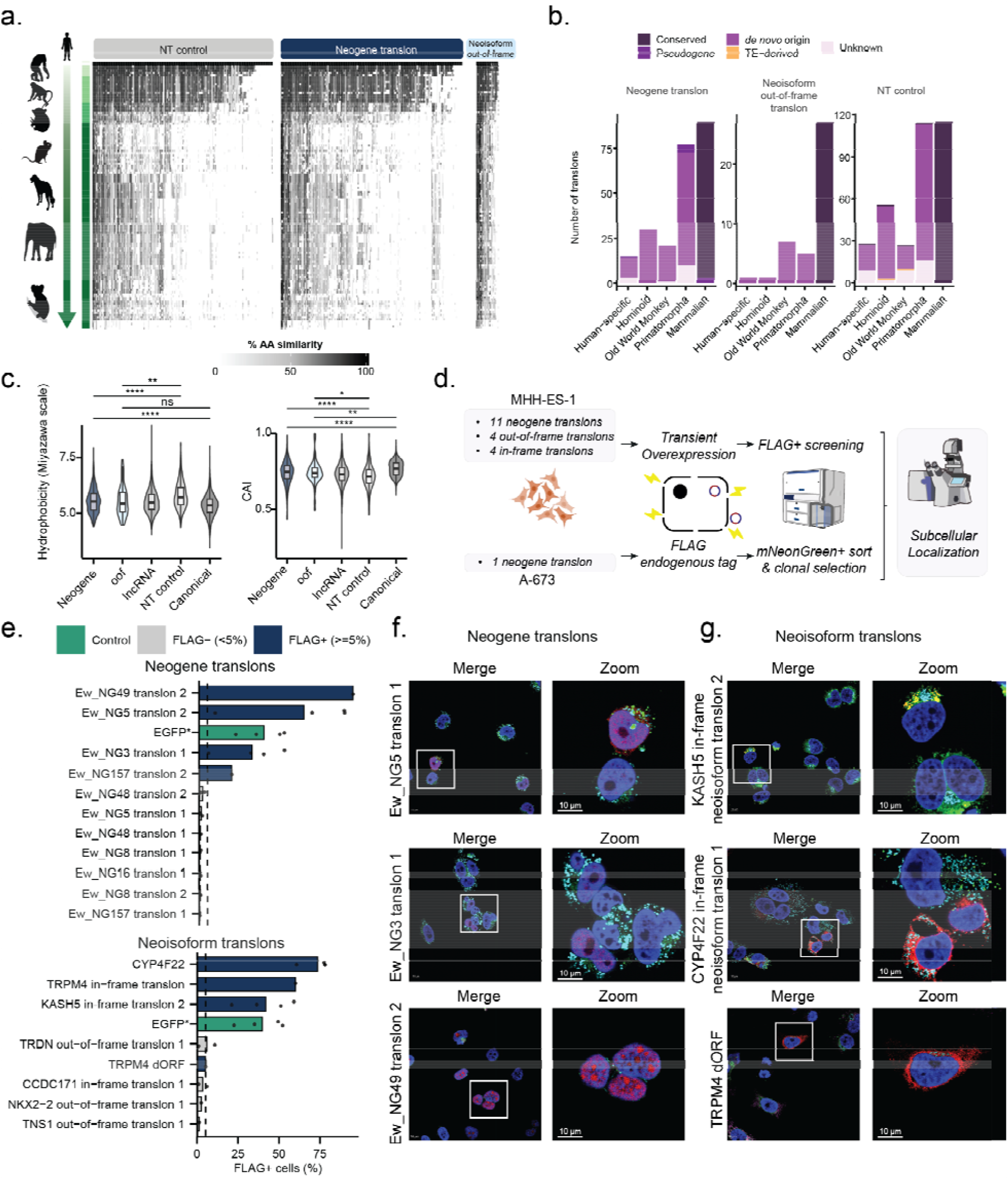
Neoproteins emerge from neutrally evolving genomic sequences, are biochemically translation-compatible and are compartment localized. **a.** Heatmap showing percentage of amino acid similarity for eac translon across species ordered by phylogenetic age, split by translon type. **b.** Number of translons classified by evolutionary mechanism of origin across phylogenetic ages, split by translon type. **c.** Hydrophobicity (Miyazawa scale) and Codon Adaptation Index (CAI) scores for the (neo)proteins encoded by the translons (oof = Out-of-frame neoisoform) compared to canonical genes, lncRNA translons, length-matched non-translated controls (NT control). Significance was tested by the Wilcoxon Rank-Sum test (n.s., p > 0.05; *p ≤ 0.05; **p ≤ 0.01; ***p ≤ 0.001; ****p ≤ 0.0001). **d.** Strategy for EwS-specific translons subcellular detection. Transient overexpression of 19 neoproteins was achieved through MHH-ES-1 cell line electroporation, FLAG-tag detection in FACS and immunofluorescenc stainings. **e.** Barplot showing the number of FLAG+ cells after transfection of overexpression constructs containing neoprotein translons, split by neotranscript type. EGFP transfection control was the same for both groups. Dots indicate separate transfection and flow cytometry experiments **f**. Merge and zoom of FLAG-Tag (red), DAPI (blue), ER (green), and Mitochondria (teal for Ew_NG3 and Ew_NG5) or Golgi (teal for Ew_NG49) immunofluorescence stainings show nuclear subcellular localization of Ew_NG5 translon 1, Ew_NG3 translon 1 and Ew_NG49 translon 2 neoproteins **g.** Merge and zoom of FLAG-Tag (red), DAPI (blue), ER (green), and Mitochondria (teal for CYP4F22) or Golgi (teal for KASH5 and TRPM4 dORF) immunofluorescence stainings show TRPM4-dORF localization to the cytoplasm, KASH5 in-frame neoisoform translon 2 (MSC-EF1_00035195_394_555) and CYP4F22 in-frame translon co-localizes to the ER.

Surprisingly, one of the most abundantly detected EwS-specific neoprotein, encoded by Ew_NG3, is an inactive pseudogene of the FSHD region 2 Family member (FRG2) gene family. This single reactivated locus produces both ancestral- and novel-sequence neoproteins. Within the eight translons, two retain detectable homology to the protein-coding FRG2 paralogs (BLASTP), while the remainder represent de novo sequences with no canonical counterpart. Multiple sequence alignment indicated the pseudogenic translons originated in the primatomorpha clade (∼70-80 mya), where it rapidly accumulated numerous disabling mutations **(Figure S10d)**. This indicates that the EWSR1::FLI1 fusion can revive expression of ancient genetic material that remained non-functional and silent for over 70 million years, until cancer-specific reactivation in EwS.

### Neoproteins exhibit intrinsic translation-compatible biochemical features

Consistent with neutral emergence, neoproteins were small (**Figure S9a-b**) and neither Kozak scores, nor protein domains, were enriched compared to NT controls (**Figure S11a-b**). Similarly, none of these features, nor the translation rates of neogenes, correlated quantitatively with evolutionary age (all R < 0.07, all p > 0.4; **Figure S11c-d**), indicating no connection between the age of the translon and its molecular characteristics. Expectedly, and in line with their cancer-restricted expression, virtually all neogene and out-of-frame neoisoform translons lacked signs of evolutionary constraints **(Figure S11e)**. Putatively functional features, such as disordered regions and signal peptides, occurred but were found at comparable frequencies in NT controls, indicating they arose by chance (**Figure S11b**). In contrast, two biochemical features were specific to neoproteins: (i) the C-terminal hydrophobicity of neoproteins was elevated relative to both canonical genes and NT controls **(Figure 6c)**, a feature identified as a signal for proteasomal degradation^55^ that could contribute to the rapid turnover of neoproteins and their enrichment in HLA-I immunopeptidomics **(Figure 4)**, and (ii) the codon adaptation index was higher in both neogenes and out-of-frame translons compared to NT controls **(Figure 6c)**.

These characteristics suggest that, by chance, some neoproteins were biochemically compatible with translation even before their transcripts were activated by fusion binding. Paradoxically, the same properties that permit transient protein existence, in particular elevated C-terminal hydrophobicity, may favor proteasomal processing and HLA-I presentation, making neoproteins inherently suited as immunotherapy targets.

### Neoproteins localize to distinct subcellular compartments

We next assessed whether neoproteins could be stably expressed and localized within cells, despite their recent evolutionary emergence and highly cancer-restricted expression. To do so, we first performed transient overexpression of 19 neoproteins **(Figure 6d, Figure S12a)**. These 19 candidates encompassed neoproteins translated from 11 neogene translons, 4 in-frame neoisoform translons, and 4 out-of-frame neoisoform translons, selected based on proteomics and immunopeptidomics detection, translation levels, patient recurrence, fusion regulation, and manual inspection of the Ribo-seq P-site profiles **(Supplementary Table S10)**. To minimize artifacts from tag-induced misfolding or aggregation, we used small C-terminal FLAG tags (8 aa). Using a flow cytometry-based readout, we were able to detect expression of 8 out of 19 neoproteins (**Figure 6e, S12d**). To independently validate this flow cytometry finding, we performed immunofluorescence to directly detect FLAG-tagged neoprotein expression as well as their subcellular localization **(Figure 6f-g, Supplementary Figure 12f)**. As shown in **Figure 6f**, Ew_NG3 and Ew_NG5 both encode detectable nuclearly-localized neoproteins, corresponding to their DeepLoc^49^ predicted nuclear localization signals. In contrast, the TRPM4-dORF was found in the cytoplasm, again matching its predicted location **(Figure 6g)**.

Interestingly, the *KASH5* in-frame translon 2 neoprotein, which still retained its TM-domain **(Figure 3e)**, was localized to the ER **(Figure 6g)**. This contrasts with canonical full-length KASH5 which typically localizes to the nuclear membrane, bringing molecular motors like dynein to the nuclear envelope^56^. Due to the loss of both EF-hand domain and signal peptide, the KASH5 in-frame neoprotein either resides in the ER or is secreted, consistent with DeepLoc prediction. Consequently, this neoprotein is likely unable to interact with its canonical interaction partners.

Additionally, we genetically engineered A-673 EwS cells using CRISPR–Cas9-mediated homology-directed repair (HDR) to knock-in a FLAG-P2A-mNeonGreen cassette at the endogenous locus of Ew_NG157, tagging its C-terminus **(Figure S12b-c)**. Ew_NG157 ranked among the top 4 highest translated neogenes and, concordantly, immunostaining of this endogenously tagged neogene revealed nuclear expression at consistently high levels, again matching DeepLoc prediction **(Figure S12h)**.

We next investigated whether stably expressed neoproteins might influence EwS cell growth. Transient overexpression of five stably expressed neoproteins in A-673 cells did not enhance proliferative capacity compared to empty-vector controls **(Figure S12e).** Similarly, knockdown of 7 neogenes (encoding 102 neoproteins) using siRNAs showed no appreciable effect on cell growth, indicating that this subset of neoproteins is unlikely to contribute to EwS cell proliferation in cell culture conditions **(Figure S13a, Supplementary Table S11)**. CRISPR interference (CRISPRi) targeting Ew_NG48 with sgRNAs directed against DNA sequences flanking EF1-bound GGAA microsatellite resulted in downregulation of Ew_NG48, consistent with a regulatory role of these GGAA repeat–bound elements in controlling Ew_NG48 expression, although this downregulation did not result in an associated phenotypic change **(Figure S13b-c, Supplementary Table S12)**. Finally, we performed shRNA-mediated inhibition of Ew_NG48 in A673/TR/shEF1 cells using 3 different IPTG-inducible shRNA constructs. Knockdown of Ew_NG48 expression did not impact cell proliferation after 72h and 6 days of IPTG treatment **(Figure S13d-f)**.

Collectively, these findings demonstrate that a subset of neoproteins can be stably expressed in cells and localize to distinct cellular compartments, including the nucleus, mitochondria, and ER, although any putative molecular roles of neoproteins in EwS remain to be determined.

## Discussion

Cancer cells express proteins that are not encoded by known ORFs, collectively referred to as the cancer “dark proteome”. However, the mechanisms that induce the dark proteome in a cancer-specific manner and the relation of dark proteome products to the oncogenic cancer cell state have largely remained elusive. EwS provides an informative case example as all EwS are driven by a single oncogenic fusion protein (EF1 in 85% of cases) that drastically remodels the cell’s gene expression landscape. In our prior work, we demonstrated how EF1 activates 26 new genes, termed neogenes^19^, but the full scope to which EF1 shapes the EwS dark proteome and the mechanisms through which it does so were not fully clear.

In our current study, we show how EF1 recurrently induces the expression of thousands of EwS-specific neoisoforms transcribed within hundreds of known protein coding genes, often through newly established intragenic transcription start sites. Using long read RNA sequencing, we also extend the set of known neogenes from 26 to 124 and provide detailed and full-length transcript maps across these usually dormant genomic sites.

Evaluating neoprotein sequences encoded by neoisoforms through Ribo-seq, we find several neoproteins partially resembling known proteins, but containing altered domain architectures or drastic N-terminal truncations that remove essential domains and change the protein’s localization. In contrast, neogenes and out-of-frame neoisoform translons produce entirely novel neoproteins not matching any known protein sequence. Particularly important for future therapeutic applications is the high recurrence of neoprotein translation as measured by Ribo-seq across 48 patient tissues and multiple genetically engineered patient-derived cell models in which we perturb EF1’s expression. Strikingly, neoprotein translation rates are on average 2-fold higher than known dark proteome constituents, such as microproteins and peptideins translated from lncRNA translons, sometimes even surpassing those of well-characterized EwS fusion targets. Our tumor profiling efforts establish neoproteins as a previously unrecognized layer of tumor-specific protein diversity and abundance.

But how could all these normally dormant protein sequences suddenly become available for transcription and translation, so that the introduction of a single fusion protein could activate them? Comparing the genomes of hundreds of species, we demonstrate that neogenes and out-of-frame translons emerged from previously non-coding sequences through a de novo mechanism in primates, similar to how other nc-translons producing dark proteome constituents evolved^24,39,57^. Under canonical evolutionary constraints, de novo genes typically exhibit low expression levels to minimize metabolic waste and proteotoxic risk^58^. The EF1 fusion bypasses this: neogene and out-of-frame neoisoform translons are expressed at strikingly high levels without any prior evolutionary refinement - in patient tumors some rival essential housekeeping genes - likely imposing a proteomic burden that gradual selection against non-beneficial proteins would normally prevent. Surprisingly, we demonstrate that the fusion can also reactivate previously dormant and disabled pseudogenes, providing the cell with both partially conserved homologues and completely novel sequences arisen de novo from the same pseudogenic loci over time. These evolutionary comparisons may also explain why rodents perform poorly as models for EwS^59^: Only 34% of the neoprotein sequences we discovered align to the mouse genome, underlining the need for patient-derived models^10,44^.

Having established the mechanisms driving neoprotein expression, we next sought to understand their putative roles in EwS oncogenesis. Through transient neoprotein overexpression and silencing experiments we could not detect measurable growth phenotypes, raising the question of their significance for EwS biology. This does not exclude a biological role: neogenes may be tolerated passenger byproducts of promiscuous fusion binding within EF1-associated transcriptional hubs^60^, may confer context-dependent advantages not captured by standard in vitro proliferation assays, or may act only through synthetic or cooperative interactions revealed upon co-depletion or co-overexpression.

Several biochemical characteristics and molecular features might imply a rapid turnover of neoproteins. For example, their elevated C-terminal hydrophobicity, lack of structure, and generally poor detectability in global proteome data align with the defective ribosomal products (DRiPs) model, wherein unstable proteins are preferentially channeled to proteasomal degradation and HLA-I presentation^55,61^. This positions neoproteins as a promising class of tumor-restricted immunotherapy targets whose inherent instability may paradoxically be advantageous, ensuring continuous antigen supply to the HLA-I pathway regardless of steady-state protein abundance. Our *in silico* simulations suggest that a neoprotein-based vaccine formulation could provide broad coverage across diverse HLA backgrounds, though experimental validation of immunogenicity and T-cell reactivity will be essential.

However, not all neoproteins are rapidly degraded. A subset shows strong evidence for stable protein expression, both through broad support in global proteomics data and stable expression of ectopically tagged constructs. Examples include the Ew_NG3 neoprotein, which is found across all our proteomics datasets, including recurrently in patients, and stably localizes to the nucleus as predicted. A particularly striking case is the *TRPM4*-dORF, an 80-amino acid neoprotein recurrently detected across all global proteomics datasets with 5 distinct peptides, yet completely absent from non-EwS tissues. Its translation appears to involve readthrough into the poly(A) tail in the absence of effective non-stop decay. That poly(A) readthrough in EwS is enriched specifically among neoisoform-bearing genes, and largely absent from non-malignant tissues, raises the intriguing possibility that fusion-driven transcriptional rewiring may alter translational fidelity beyond simply activating new transcripts. *EWSR1::FLI1* is known to regulate alternative splicing^62,63^ and mRNA decay^64^. Whether the poly(A) readthrough we observe reflects a loss of normal EWSR1-mediated RNA processing at affected loci, or instead results from altered deadenylation or 3′ UTR architecture at fusion target genes, remains to be determined.

Central questions in the emerging field of dark proteome research include whether a cancer-specific dark proteome exists, how it is induced, and whether its products have functional or therapeutic relevance. Our findings in EwS address several of these questions. We show that a single oncogenic fusion protein is sufficient to activate a large dark proteome by rewiring the transcriptional landscape at normally silent genomic loci, coupling dark proteome induction directly to the primary oncogenic event rather than to stochastic transcriptional noise. Whether the resulting neoproteins are functional remains less clear; most appear to be rapidly turned over, and individual perturbation did not yield measurable growth phenotypes. Yet their collective abundance and presence on HLA-I molecules suggest the dark proteome is not biologically inert, even when individual products lack detectable function. More broadly, the human genome appears to harbor a reservoir of sequences compatible with translation that remain silent under normal conditions. EwS, in which a single fusion protein inadvertently accesses this reservoir, serves as a natural experiment that illuminates the genome’s hidden protein-coding capacity, with relevance for tumor-specific immunotherapy and for our understanding of how new proteins arise.

Our dataset, representing the deepest cancer translatome generated to date, comprises over 4 billion RPFs from 48 patient samples matched by proteome and transcriptome data, provides a comprehensive atlas of 124 neogenes and over 2,000 neoisoforms. All data and annotations are publicly available as a resource for future mechanistic and immunotherapeutic studies of fusion-driven transcriptome expansion.

## Methods

### Resource availability

Ribosome profiling data and RNA-seq for 1 patient from the Princess Máxima Center for Pediatric Oncology biobank was deposited at the European Genome-phenome Archive (EGA) with accession number EGAS00001008485. All other ribosome profiling data and newly generated patient short-read RNA-seq data was deposited at the German Human Genome-phenome Archive (GHGA) with accession number GHGAS26808925086395. Long-read RNA-seq data was deposited at GEO with accession number GSE338815. Single-cell Ribo-seq raw data was deposited at GEO with accession number GSE337865. Newly generated LC-MS/MS proteomics raw files for dataset 1 and the search output files have been deposited to the ProteomeXchange Consortium via the PRIDE partner repository under the following identifier: PXD081387. Newly generated LC-MS/MS proteomics raw files for dataset 3 and the search output files have been deposited to the ProteomeXchange Consortium via the PRIDE partner repository under the following identifier: PXD079451. Newly generated immunopeptidomics raw files for dataset 4 and the search output files have been deposited to the ProteomeXchange Consortium via the PRIDE partner repository under the following identifier: PXD081499.

### Code availability

All custom code generated for this study is available on the following Github pages: https://github.com/lbroeil/Broeils-et-al-2026-Ewing-Sarcoma, https://github.com/JillPilet/Dark_proteome_ewing_sarcoma https://github.com/mkkoshkina/neoprotein_landscape_of_EwS_data_analysis/tree/main

### Cell lines and reagents

All TR/shEF1 and TR/shERG EwS cell lines were cultured in Roswell Park Memorial Institute (RPMI) 1640 medium with Glutamax (Gibco), supplemented by 10% (tested to be dox-free) fetal bovine serum (FBS)(Gibco), 1% penicillin & streptomycin (Gibco) and incubated at 37 °C, 5% CO2 in culture flasks (Thermo Fisher). The WT A-673 cell line was cultured in DMEM supplemented with 10% FBS (Gibco) and 1% penicillin & streptomycin (Gibco). Ewima1 cells were cultured in Alpha MEM, 10% MSC-specific FBS supplemented with 10mM L-Glutamine and 1% penicillin & streptomycin (Gibco). TC-71, EW-16 and EW-7 cell lines were cultured in RPMI (Gibco), supplemented with 10% FBS (Gibco) and 1% penicillin & streptomycin (Gibco). TC-71 and EW-16 were cultured in pre-coated flasks with collagen from bovine skin (C4243; Sigma-Aldrich, St. Louis, MO, USA).

EWIma1 clones were generated by engineering an *EWSR1::FLI1* translocation in MSCs as described previously^35^. The TR/shEF1 and TR/shERG EwS cell lines were established as described previously^10^. In short, cells were transduced with a pLKO-Tet_on all-in-one lentiviral vector containing a Tetracycline repressor gene that is constitutively active, and a shRNA targeting the RNA of the respective driver fusion-protein. To enhance growth, RD-ES/TR/shEF1 cells were cultured in flasks pre-coated with 100 µg/cm^2^ gelatine from bovine-skin (Sigma-Aldrich) for 1h. Cells were passaged twice a week at differing densities, depending on the cell line. All cells routinely tested negative for mycoplasma infection and STR fingerprinting was done every two months to confirm cell line identity.

### Primary patient material

Archival fresh-frozen primary EwS patient samples (n = 54) from biopsies or resections were provided by the Department of Orthopedics, Heidelberg University Hospital (Heidelberg, Germany) or kindly provided as left over INFORM^65–68^ material. From the biobank of the Princess Máxima Center for Pediatric Oncology, one sample was received. Samples used for Ribo-seq and RNA-seq were kept at –80 °C, then powdered using a tissue pulverizer. All samples were harvested, processed and used according to approved biobank proposals and signed informed consent forms.

### Cell line fusion knockdown experiments

To induce the knockdown of the fusion, the cell lines with a short hairpin RNA against the respective fusion driver (A-673/TR/shEF1, MHH-ES-1/TR/shEF1, SK-N-MC/TR/shEF1, RD-ES/TR/shEF1, TC-106/TR/shERG) or shControl (A-673/TR/shControl) were treated with 1 µg/mL doxycycline hyclate (dox) (Sigma-Aldrich) per mL of cell culture medium for 96 h. Dox was refreshed every 48 h.

### Lysis and cell harvest for RNA-seq and Ribo-seq

Powdered patient material was lysed in 600–1,000 µL of 1X lysis buffer (20 mM Tris-Cl pH 7.4 (Sigma-Aldrich), 150 mM NaCl (Fisher Scientific), 5 mM MgCl_2_ (Fisher Scientific), 1% Triton X-100 (Sigma-Aldrich), 0.1% Igepal CA-630 (Sigma-Aldrich), 1 mM DTT (Lucigen), 10 U/mL RNase-free DNase 1 (Lucigen), 0.1 mg/mL cycloheximide (Sigma-Aldrich) in Nuclease-Free Water (Sigma-Aldrich)), depending on weight of the material. Lysates were kept on ice for 10 min, being vortexed every two minutes. Lysates were then centrifuged at 20,000 g for 10 min, after which cleared lysates were split into 200 µL fractions. For the cell lines, 10–20 million cells were harvested after 96 h dox treatment by aspirating cell culture medium, then washed with ice-cold PBS supplemented with 0.1 ng/mL of cycloheximide (Sigma-Aldrich), while the plate is kept on ice. The cells are then lysed in plate with 1X lysis buffer and processed with an equivalent protocol as the patient material.

### RNA isolation for short-read Illumina sequencing

RNA was extracted from lysates by TRIzol LS (Invitrogen) – Chloroform (Sigma-Aldrich) extraction, followed by purification using the RNA Clean and Concentrator-5 kit (Zymo) according to manufacturer instructions. The isolated RNA was then treated with 1 U/µL DNase I (Lucigen) for 10 min at 37 °C, followed by a purification step using the RNA Clean and Concentrator-5 kit (Zymo). RNA concentration was measured using the NanoDrop (Thermo Scientific) and RNA integrity (RIN) value was determined by the Bioanalyzer 2100 (Agilent) using the RNA 6000 Nano kit (Agilent).

### Short-read Illumina RNA-sequencing

Isolated RNA with a RIN value above 6 was poly(A) enriched by Azenta Life Sciences and subsequently sequenced according to their RNA-seq NGS services.

### Transcriptome assembly – short read

Raw reads were trimmed using fastp^69^ and then mapped to the human genome (hg38, Ensembl v102) using STAR^70^ with the following parameters: *--outFilterType BySJout --outSAMunmapped Within -- outSAMattributes NH HI AS nM NM MD jM jI MC ch --outSAMstrandField intronMotif --outSAMtype BAM Unsorted --outFilterMismatchNmax 6 --alignSJoverhangMin 10 --outFilterMultimapNmax 10 -- outFilterScoreMinOverLread 0.75*.

To create the short read transcriptome, 133 EwS RNA-seq samples from four different sources (the Princess Máxima Center for Pediatric Oncology repository (n = 51), the St. Jude Cloud (n = 37), produced and kindly provided by the INFORM programme^65–68^ (n = 7) and this study (n = 38)) were processed with StringTie^71^ using the Ensembl v102 as a reference and with the following parameters: *-M 0.95 -a 9 -m 100 -f 0.05 -g 40 -c 10 -j 2 -s 99999*. GFFcompare^72^ was then used to merge all sample-specific GTF files into one GTF file, which was filtered to only include transcripts occurring in a minimum of 3 samples and classified as transcript class *k, o, x, I, y, and u* as defined by GFFcompare^72^.

### Short read RNA-seq quantification

To quantify using Salmon^73^ v1.8, first an index for the EwS transcriptome was created. After trimming using fastp^69^ v0.23.4, reads were quantified on the transcript level using Salmon using standard parameters. Quantification files were loaded into R as a DESeq2 object using the tximport package, with *countsFromAbundance* set to *scaledTPM* and transcripts being summarized into gene-level counts.

### Identification of neogenes in short-read RNA-seq data

To define EwS-specific neogenes, we quantified the expression of the EwS-transcriptome across three comprehensive reference datasets using Salmon v1.8.0 and performed differential expression analysis using DESeq2 v1.44. The reference datasets comprised: (1) the GTEx dataset, spanning 54 non-malignant tissue types consolidated into 23 groups^74^; (2) the Evo-Devo atlas, containing developmental transcriptomes from 7 tissue types across multiple timepoints from 4 weeks post-conception through adulthood^36^; and (3) a pediatric cancer cohort of RNA-seq from ten non-EwS tumor types (AML, ATRT, B-ALL, EPN, MBL, NBL, OS, RMS, T-ALL, and WT) from the St Jude Cloud and Princess Máxima Center for Pediatric Oncology. For each dataset, we performed pairwise differential expression analyses comparing EwS samples against: (i) each tissue group from GTEx; (ii) each tissue type from Evo-Devo, stratified by pre- and post-birth timepoints; and (iii) each distinct tumor type from the pediatric cancer cohort.

LncRNA, pseudogenes and unannotated genes were classified as neogenes if they had a min LFC > 1, a mean LFC > 3 and an adjusted p-value of < 0.005 across all comparisons, with the exception of the testis tissue groups in (i) and (ii). In addition, we required a min LFC > 1 in the RNA-seq quantification of at least 1 EwS TR/shETS cell lines after performing differential expression analyses using DESeq2 v1.44. Lastly, we required the neogene TSS to be within a TAD containing EF1 bound GGAA microsatellites defined previously^75^.

### Nanopore and PacBio long-read sequencing

RNA was isolated from A-673, TC-71, EW-7, EW-16 and Ewima1 cell lines using TRIzol reagent (Invitrogen) and chloroform. After elution, RNA was split for Nanopore and PacBio processing as followed:

- 70% of RNA was treated with TURBO DNase (Ambion, Life Technologies #AM2238) to avoid DNA contamination as specified by the supplier. Messenger RNAs were then selected through Dynabeads poly(A) selection (Invitrogen #61006). Libraries were then prepared from 500ng poly(A)+ RNA using the Direct RNA sequencing protocol (SQK-RNA002, Oxford Nanopore technologies) and samples were sequenced on an Oxford Nanopore Technology MinIon sequencer (Mk1C). Sequencing was performed for 48-72 h.

- The remaining 30% of RNA transcripts were purified using Nucleospin RNA (Macherey-Nagel #740955.50) with rDNase treatment. All libraries were prepared with SMRTbell Express Template Prep Kit2.0 and samples were sequenced on PacBio Sequel II sequencer.

### Nanopore assembly

Base calling was performed with guppy v5.0.16 to generate FASTQ files from Nanopore output FAST5 files. FLAIR^76^ v1.6.3 was used to align and generate transcriptome. The module “align” was applied with minimap2^77^ v 2.17 on genome reference hg19. We also used the modules “correct” and “collapse” to take into account short reads sequencing (junctions detected by STAR on Illumina RNA-seq) and GENCODE v19 annotation.

### PacBio assembly

Isoforms clustering was performed on HiFi reads FASTQ files followed by mapping on reference genome hg19 using pbmm2 v1.0.0. Aligned isoforms bam files were then compared with reference transcriptome v19 and annotated with SQANTI^78^ (Isoseq3) tool.

Of note, although the initial assemblies of Nanopore and PacBio data were performed on hg19, all integration with short-read sequencing and Ribo-seq data were performed on hg38 through batch coordinate liftOver and GFFcompare^72^.

### Identification of neoisoforms and neogenes in long-read RNA-seq

The detailed pipeline to identify neoisoforms and neogenes in long read sequencing is available **Figure S1a**. After Nanopore and PacBio transcripts assembly, all transcripts from the 2 techniques were merged per cell line. We used HTSeq-count with the new assembly to count the number of reads per gene and transcript in the short-read sequencing. We then performed a multi-step filtering to identify highly EwS-specific sequences for each cell line. First, we kept only highly expressed genes (>15TPM with Illumina short-read sequencing). We then selected isoforms:

- With a EWSR1::FLI1 binding at a GGAA microsatellite in the same topologically associated domain (TAD). To do so, we annotated TADs from Showpnil et al. Nucleic acid research, 2022.
- Overlapping an active mark (H3K27ac or H3K4me3 ChIPseq peak).
- Downregulated upon 72 h of EF1 knockdown (log2FoldChange _WT vs KD_>0.5 in A-673 or TC-71.
- Not expressed in 6 MSCs cell lines (transcript expression ≤0.03TPM).

We then merged all transcripts from the 5 cell lines (A-673, TC-71, EW-7, EW-16 and Ewima1 clone) using GFFcompare^72^. We only kept isoforms with high-expression levels at the transcript level (Median >1.5TPM in a cohort of 33 EwS cell lines or >1.2 in Ewima1 model or at least 10 reads in Nanopore sequencing). Two classes of transcripts were then made, one “neoisoform” class characterized by a match in GENCODE reference transcriptome and a “Neogene” class with no match in GENCODE.

To identify highly EwS specific neogenes we filtered Neogenes with a mean expression <5 in non-EwS TCGA and GTEx datasets and a fold change in EwS vs non EwS datasets >15 in the Illumina or Nanopore dataset. Finally, to select EwS-specific neoisoforms, we removed “full-splice match” isoforms annotated with SQANTI leading to 2,244 isoforms of 252 genes (**Supplementary Table S6**). While 54% of canonical genes with neoisoforms were detected by both long-read technologies (135 out of 252), PacBio sequencing provided a greater depth of neoisoform detection (1,715 vs 391 neoisoforms identified in Nanopore sequencing and 138 identified in both technologies), with most neoisoforms detected across multiple cell lines **(Figure S1b-e)**. Apart from isoforms with neoexons and intragenic TSS isoforms, 1,484 non-canonical neoisoforms resulted from multiple alternative splicing events. These isoforms displayed at least one novel splice site (n = 518), a combination of known splice sites (n = 242), intron retention (n = 192), combination of known junctions (n = 180), 3′ fragment (n = 192), mono-exon (n = 88), 5′ fragment (n = 46), multi-exon (n = 17), mono-exon by intron retention (n = 8) and internal fragment (n = 1) **(Figure S5b-c)**.

### RNA-seq quantification of neoisoforms

Neoisoforms quantification can be challenging for sequences less than 25 bp length due to the homology with canonical sequences flanking the EwS-specific sequence. To quantify the neoisoforms expression levels, we extracted these EwS-specific sequences in each isoform by sequence comparison to GENCODE. We then excluded sequences shorter than 25 bp as this is the minimum length of our minimizer indices and we filtered EwS-specific sequences with mean expression <1 in non-EwS datasets, FoldChange expression in Illumina or Nanopore >15 and a minimum expression in Illumina or Nanopore in EwS >10. This extensive filtering yielded 594 EwS-specific isoforms with neoexons for which the expression could be estimated in TCGA and GTEX datasets using NEEDLE quantitative minimizer based approach^79^.

### Neogene and neoisoform expression estimates

Neogene and neoisoform expression in TCGA, GTEx and target custom Ewing Sarcoma index was estimated as described previously^80^. Briefly, we indexed each cohort with Needle v1.0.1^79^ (needle ibf -w 25 -k 20 --cutoff 2 -l 10 -n 5 -f 0.05), which couples an interleaved Bloom filter (IBF)^81,82^ with minimizer-based sequence reduction^83^: for each sample, the IBF stores the presence of every minimizer over sliding windows. For bulk data, an index was built per project (33 TCGA, 54 GTEx) from concatenated paired-end fastq files. For neogenes, full length sequences were queried. For neoisoforms, only the EwS specific portion of the transcript sequence was queried. Query sequences were grouped into batches of fewer than 100 million nucleotides and searched with needle estimate (Needle v1.0.3), which returns for each sample the median of the minimizer counts of the query. Because this median does not depend on the number of minimizers extracted, it is independent of query length, so length normalization is inherently applied. The resulting expression tables were converted into sparse matrices (scipy.sparse.csr_matrix; SciPy v1.15.1^84^) and encapsulated into AnnData objects (anndata v0.11.3). Estimates were then divided by the total read count of the sample and multiplied by 10 to obtain depth-normalized expression levels, applying the TPM normalization principle^85^ to make values comparable across samples.

### Annotation of genes with intragenic TSS isoforms

Isoforms were annotated as Intragenic TSS isoforms when TSS colocalized with a GGAA microsatellite located in the genomic sequence of a canonical gene and when the transcript was shorter than the main canonical transcript. The annotation of intragenic TSS isoforms was then performed at the gene level through manual curation to reach 32 genes.

### Merging of short-read and long-read transcriptomes

The gtf file containing the long-read assembled neogenes and neoisoforms was first converted to hg38 using UCSC liftover. This file was then merged with the short-read transcriptome assembly gtf using GFFcompare^72^ using the GENCODE reference v36. After merging, in total 124 neogenes were assigned. Numbering did exceed 124 because some neogenes were filtered out later in the process.

### CAGE-data analysis

CAGE-data from SK-N-MC and HepG2 cell lines was obtained from FANTOM^37,38^. BED files were converted to BigWig format for genomic visualization. CAGE signal intensity was plotted around the transcription start sites (TSSs) of neoisoforms, spanning ±3 kb, using the computeMatrix and plotHeatmap functions from the deepTools Python package. The matrix was generated using a bin size of 50 bp. For plotting, samples were ordered according to CAGE signal intensity in the SK-N-MC cell line, both on the sense and antisense strands.

### Ribo-seq

Ribosome profiling was performed as described previously^27^. In short, lysates were digested with RNase I after which the RNA was isolated. We then depleted the non-mRNA molecules using antisense probes (RiboPOOL, siTOOLS) and performed size selection for expected RPF length (28-30bp) using gel electrophoresis. Sequencing adapters were ligated to the RPFs, followed by reverse transcription, gel electrophoresis size selection and circularization. After PCR amplification, a final gel electrophoresis step to select for RPF-containing libraries, and clean-up, final libraries were combined in equimolar concentrations and sequenced on the NextSeq2000 P3 flowcell (Illumina), the NovaSeq 6000 S4 flow cell (Illumina) or the NovaSeq X 10B flowcell (Illumina), with 50 cycles for read 1 and 6 cycles for the barcode.

### Ribo-seq read processing, alignment and QC

Raw reads were trimmed using Trimgalore^86^ v0.6.6 using standard parameters, except *–length 25*. Trimmed FASTQs were then filtered for rRNA, tRNA, snRNA, snoRNA, and mtRNA by mapping to an in-house index using Bowtie2^87^ v2.4.2 with standard parameters, except *–L 25.* Reads that did not align to any of the non-mRNA sequences were then mapped to the human genome (hg38) using STAR^70^ v2.7.8a with the EwS-transcriptome GTF file as a reference, a 29nt STAR index and the following parameters: *--runDirPerm All_RWX --twopassMode Basic --outFilterMismatchNmax 2 --outFilterMultimapNmax 20 -- outSAMattributes All --outSAMtype BAM SortedByCoordinate --quantMode GeneCounts -- limitOutSJcollapsed 10000000 --limitIObufferSize=300000000 --outFilterType BySJout -- alignSJoverhangMin 1000*. Aligned BAM files were used as input for RiboseQC^88^, which generated periodicity scores, codon usage and other QC stats.

### Translon-calling, reannotation and filtering

To maximize Ribo-seq coverage, we implemented a sample-merging strategy as it has been shown to improve translon identification^89^. First, *`for_ORFquant`* files generated by RiboseQC were merged while preserving sample-specific P-site cutoff determinations. This merged file served as input for ORFquant^90^ v1.0, which was executed using default parameters.

For PRICE^41^ analysis, trimmed and filtered FASTQ files were mapped using STAR v2.7.8a with parameters identical to those used for ORFquant, apart from using `--alignEndsType EndToEnd`. The resulting BAM files were merged using samtools v1.12, and PRICE from the gedi v1.0.6 toolkit was run with default parameters. The *`orfs.cit`* output was converted to BED format for subsequent processing. Output files from both translon-calling tools were imported into R and standardized to compatible formats.

Each identified translon was systematically classified based on its genomic location, reading frame, and host gene biotype. translons within protein-coding genes were categorized as canonical CDS isoforms (any in-frame CDS overlap), upstream ORFs (uORFs), upstream overlapping ORFs (uoORFs), internal ORFs (intORFs), downstream overlapping ORFs (doORFs), or downstream ORFs (dORFs), based on their position relative to the annotated CDS start and stop codons. Translons in non-coding regions were classified as lncRNA, pseudogene, or unannotated gene translons based on host gene annotation.

To accurately quantify translation levels for each translon, we implemented a P-site-based measurement approach. First, we generated BED files containing all genomic P-site positions for each translon identified by our translon-calling pipeline. These files were then intersected with sample-specific BED files containing all P-site coordinates (as determined by RiboseQC) using Bedtools^91^ intersect v2.31.0 with parameters *`-wa -wb -header -f 1.00 -s`*.

From these intersections, we calculated the number of in-frame P-sites for each translon in each sample, based on the reading frame established by the start codon. To normalize these counts, we developed a P-site Per Million (PPM) metric, analogous to TPM for RNA-seq data, but specifically designed for ribosome-protected fragments. The PPM calculation involved:

1. Determining P-sites per kilobase for each translon by normalizing raw in-frame P-site counts to translon length
2. Calculating a sample-specific scaling factor by dividing the sum of all P-sites per kilobase values by 1,000,000
3. Dividing each translon’s P-sites per kilobase value by this scaling factor

This normalization approach enables direct comparison of translation efficiency across translons of different lengths and between samples with varying sequencing depths. To only keep recurrently expressed translons and avoid sample-specific translation, we only kept translons in our list that have a PPM > 1 in at least 10 patient samples (out of 48).

For Ribo-seq data generated from EwS cell lines, we applied the same P-site quantification methodology described above. However, to maintain consistency and enable direct comparison with patient data, we restricted our analysis to the translon catalog previously established in patient samples. No additional translon-calling or filtering was performed on the cell line data; instead, we quantified translation of the patient-derived translon set across all cell line conditions. This approach ensured that translational changes observed after fusion protein knockdown were assessed using a uniform set of translons, facilitating robust comparative analyses between patient samples and experimental cell line models.

### Single-cell Ribo-seq library preparation

Single-cell ribosome profiling libraries were prepared using scRibo-seq^48^ that was modified to use RNase I instead of micrococcal nuclease. After sorting, 50 nL of RNase I (0.0195 U/µL; ThermoFisher) was dispensed to each well and incubated at 37 °C for 30 min. All remaining steps of nuclease inactivation, small RNA library construction, and purification were performed as described in scRibo-seq^48^. Single-cell ribo-seq libraries were sequenced using v1.0 chemistry on a NovaSeq X (Illumina; NovaSeqXSeries Control Software version 1.3.0.39308; bcl2fastq v2.20.0.422), with 150 cycles for read 1, 150 cycles for read 2, 6 cycles for the i7 index read (plate barcode), and 10 cycles for the i5 index read (cell barcode).

### scRibo-seq - read processing, alignment and quantification

Ribo-seq raw reads were processed, aligned, and quantified as described in scRibo-seq^48^. Reads were aligned to hg38 (GRCh38.p14), using GENCODE v49 annotations for metagene and quality control analyses and custom GTF with non-canonical translon annotations for gene translation quantifications.

### GGAA microsatellite identification and TSS assignment

GGAA microsatellites were identified as sequences containing a minimum of four consecutive GGAA repeats using the FIMO tool^92^. The TSSs were determined as follows: for annotated genes, TSSs were extracted from the hg38 Ensembl v102 GTF file; for neogenes, the TSS from the most recurrently expressed isoform was selected; for neoisoforms, isoforms with TSSs within 50 bp (intragenic TSS type) or 250 bp (other neoisoforms) were grouped and represented by a single TSS.

ChIP-seq peaks for EWRS1::ETS binding in EwS cell lines were obtained from previously published data^10^ and converted to hg38 using UCSC liftover. BigWig files for H3K27ac and H3K4me3 from the same study were first converted to BedGraph using UCSC bigWigToBedGraph, lifted over to hg38, then converted to peaks for each cell line using MACS2^93^ bdgpeakcall with -l 200 and -g 400. Peaks were then merged and kept if they were called in a minimum of 3 cell lines.

Distance to nearest GGAA microsatellite, EWSR1::ETS, H3K27ac and H3K4me3 was calculated using the GenomicRanges package^94^. Difference in distribution was tested using the Kolmogorov-Smirnov test. P-values adjusted using Benjamini-Hochberg correction.

### Overlap analysis with GENCODE ncORF set

To determine overlap with the latest GENCODE ncORF set, the comprehensive ncORF catalog was obtained^42^. From the starts and stops coordinates, each p0 site was determined for each ncORF. These p0 site genomic coordinates were then compared to all p0 site genomic coordinates from the EwS translon table containing all nc-translons. If there was any in-frame overlap of these p0 sites, translations were counted as overlapping.

### Analysis of differential expression after dox-induction of TR/shEF1 or shERG cell lines

For RNA-seq data, Salmon^73^ quantification files were loaded into R as a DESeq2 object using the tximport package, with *countsFromAbundance* set to *scaledTPM* and transcripts being summarized into gene-level counts. For Ribo-seq data, in-frame P-site matrices were loaded into R as a DESeq2 object after being summarized to gene-level P-site counts. For protein-coding genes, CDSs on MANE transcripts were taken for Ribo-seq quantification. DESeq2 was then used to to perform differential gene expression separately within each cell line, where a | log2FC > 1 | and p < 0.05 was considered a significant change.

### Protein structure predictions and visualization

KASH5 protein structure predictions were downloaded from Uniprot (https://www.uniprot.org/uniprotkb/) and Alphafold (https://alphafold.ebi.ac.uk/) databases. Protein structure predictions for KASH5 EwS-specific translons were obtained with Alphafold 3^95^ and visualized using The PyMOL Molecular Graphics System, Version 3.1.6.1, Schrödinger, LLC.

### Translon characteristics analyses

The InterProScan web service was used to scan all neogene protein FASTA sequences against the InterPro protein signature database^51^. DNA and amino acid sequences were extracted for each translon and analyzed for five molecular features. GC content was calculated using the Biostrings package. Codon Adaptation Index (CAI) was imputed with the cubar package^96^. C-terminal hydrophobicity was determined using the peptides package on the final 30 amino acids (or entire sequence for proteins <30 aa). Kozak scores were calculated by isolating a 15-nt window around the start codon (excluding the start codon itself to avoid penalties for near-cognate starts) and comparing to a position weight matrix derived from all canonical protein-coding genes using Biostrings. Intrinsic disorder scores were computed with IUPRED3^97^.

For comparison datasets, MANE transcripts were selected to represent each canonical protein-coding gene, avoiding redundancy from multiple isoforms. The lncRNA translon set comprised non-canonical translons called on genes annotated as lncRNAs. Non-translated control sets were generated using Gettranslon from EMBOSS^98^ to identify all theoretical translons on neogene transcripts. These theoretical translons were filtered to match the length distribution of neogene translons (NT ctrl). To control for potential selection bias, we generated an additional control set containing 10-fold more length-matched translons (NT ctrl 10x).

### Translon phylogenetic analyses

To determine evolutionary origin using the 120 mammal alignment^99^, we developed an in-house script based on a previously published pipeline^24^. In short, we used maf_parse from PHAST^100^ to extract the corresponding alignments for each translon based on the hg38 genomic coordinates, only keeping alignments that had a minimum of 90% coverage (including annotated gaps). These extracted sequences were then aligned using PRANK v170427^101^ to reconstruct ancestral sequences. Using this, we could define in which species an translon was conserved. We considered a translon to be conserved if the “ATG” was present, the reading frame was intact, and the translon length was >= 70% of the original length. If a translon was present in at least 1 species in a species group, we considered it that whole group, e.g. if a translon was present in tarsiers of the primatomorpha group, we considered an translon intact in the primatomorpha group. In parallel, BLASTP v2.5.0^102^ was performed using the swissprot database with a cutoff value of P < 10^-5^.

Using these data, we defined the different evolutionary origins as follows:

- If a translon originated later than the syntenic region AND it has no BLASTP results, we considered it *de novo*.
- If a translon originated later than the syntenic region AND it had significant BLASTP results, we considered it a pseudogene of its BLASTP match.
- If a translon had BLASTP matches to TE-elements, we considered it to be TE-derived
- IF a translon had significant BLASTP results or had an intact translon in the mammalian group, we considered it Conserved.

Using the multiple alignments for each translon sequence, we ran PhyloCSF^103^ using standard settings.

To visualize individual translon evolution trajectories, multiple alignment files were loaded into Jalview^104^. Loss of ATG was colored in red, premature stop codon was colored in orange and indels resulting in a frameshift are colored in yellow.

### Proteomics data analysis – tryptic global proteomics (dataset 1)

LC-MS/MS spectra were searched using FragPipe v23.1^105^ (MSFragger v4.3, Philosopher v5.1.2, Percolator v3.7.1, IonQuant v1.11.11) against a custom FASTA database combining the SwissProt Human Reference Proteome (42,520 entries, date of retrieval - 2024.10.29) with a database of 49,343 EwS-specific non-canonical translon sequences (described above), totaling 183,742 entries after addition of reversed decoy sequences and common contaminants within FragPipe.

Database searches were performed with strict trypsin as the digestion enzyme (cleavage C-terminal to K and R), allowing up to 2 missed cleavages and peptide lengths between 7 and 50 amino acids. Precursor mass tolerance was set to ±20 ppm and fragment ion mass tolerance to 0.02 Da, using b- and y-ion series. Cysteine carbamidomethylation (+57.021 Da) was set as a fixed modification. Variable modifications included methionine oxidation (+15.995 Da) and N-terminal protein acetylation (+42.011 Da). PSM validation was performed using Percolator v3.7.1 with target-decoy competition (flags: --only-psms --no-terminate --post-processing-tdc). PSM and protein-level FDR were controlled at 1% using group-specific FDR estimation in Philosopher, with PSMs grouped by protein existence (PE) level encoded in the FASTA headers — canonical UniProt entries (PE=1) and non-canonical translon-derived sequences (PE=2) were evaluated as separate groups, preventing FDR inflation from the larger canonical database. Peptide spectrum matches were rescored using MSBooster v1.3.17 with DIA-NN-based retention time and spectral intensity predictions.

For quantification, spectral counts were exported from Philosopher and Normalized Spectral Abundance Factors (NSAF) were calculated per protein as the spectral count divided by protein length, normalized by the sum of all SC/L values within each sample. Values were subsequently log-transformed as log10(NSAF + 10□□) + 5 for visualization and downstream analysis. For visualization and downstream tasks only proteins that were identified as “lead” by parsimony algorithm based on Occam’s razor principle and had no SwissProt entries in the “Indistinguishable proteins” in FragPipe results were kept.

### Proteomics data analysis - Multienzyme global proteomics (dataset 2)

Raw data from the A-673 deep multi-enzyme proteomics dataset^33^ were searched against the same custom FASTA database described above using FragPipe v23.1^105^ (MSFragger v4.3, Philosopher v5.1.2, Percolator v3.7.1, IonQuant v1.11.11). Spectra were searched separately for each of eight digestion enzymes: trypsin (cleavage C-terminal to K/R, no cleavage before P), Arg-C (cleavage C-terminal to R, no cleavage before P), Asp-N (cleavage N-terminal to D/E), chymotrypsin (cleavage C-terminal to F/L/W/Y, no cleavage before P), Glu-C (cleavage C-terminal to D/E), Lys-C (cleavage C-terminal to K), Lys-N (cleavage N-terminal to K), and Proalanase (cleavage C-terminal to A/P). All enzymes allowed up to 1 missed cleavage and required fully enzymatic termini. Peptide lengths were restricted to 7–50 amino acids and precursor charge states to 2–4.

All other search parameters matched the tryptic global proteomics workflow described above, with the following exceptions: fragment ion mass tolerance was set to 20 ppm (data acquired with ion mobility-enabled instrumentation), the top 500 peaks per spectrum were used for matching, and isotope errors of 0/1/2 were considered. PSM and protein-level FDR were controlled at 1% using group-specific FDR estimation. NSAF was calculated independently for each enzyme digest as described above, and results were merged at the detection level across all eight enzymes, retaining a protein as detected if identified in at least one enzyme condition. For visualization and downstream tasks only proteins that were identified as “lead” by parsimony algorithm based on Occam’s razor principle and had no SwissProt entries in the “Indistinguishable proteins” in FragPipe results were kept.

### Proteomics data analysis – global proteomics of patient tumor samples (dataset 3)

Fresh-frozen EwS patient tumor samples were prepared using a single-sample workflow for joint metabolomic and proteomic analysis^106^. Briefly, biphasic 75% EtOH/MTBE extraction was performed, and the resulting protein pellets were resuspended in SDS lysis buffer (4% SDS, 100 mM ammonium bicarbonate, pH 8.5) and subjected to AFA ultrasonication (Covaris LE220R-plus). Protein concentrations were determined using a BSA assay (Thermo Fisher). 12 µL of lysate per sample was processed using the automated SP3 protocol on a Bravo liquid handling system (Agilent Technologies), including protein clean-up, reduction and alkylation (10 mM TCEP, 40 mM CAA), and trypsin digestion (1:20 enzyme:protein ratio)^107^. 200 ng of peptides per sample were injected onto a timsTOF Pro mass spectrometer (Bruker Daltonics) coupled to an Easy nLC 1200 system (Thermo Scientific) with an 80 min gradient at 300 nL/min. Data were acquired in DIA-PASEF mode over a mass range of 100–1,700 m/z and an ion mobility range of 0.65–1.42 V·s/cm².

Raw files were analyzed using DIA-NN 2.2.0 in library-free mode against the same custom FASTA database described above. Trypsin/P was used as the digestion enzyme with up to 2 missed cleavages, peptide lengths of 7–30 amino acids, and precursor charge states of 1–4. Cysteine carbamidomethylation was set as a fixed modification; methionine oxidation and N-terminal methionine excision were included as variable modifications (maximum 1 per peptide). Deep learning-based prediction of spectra, retention times, and ion mobilities was enabled. Match-between-runs was enabled with RT-dependent cross-run normalisation. Precursor FDR was set to 1%. As DIA-NN does not support group-specific FDR, FDR was controlled globally across all database entries. Peptide sequences matching annotated canonical proteins were discarded. Protein group intensities were median-normalized across samples and log-transformed for visualization and downstream analysis.

### Proteomics data analysis – HLA-I immunopeptidomics (dataset 4)

HLA-I-associated peptides were isolated from five EwS cell lines (A-673, EW-7, EW-16, STA-ET-1, TC-71) and five patient-derived xenografts (IC-pPDX-5, -8, -141, -164, -179) as previously described^34^. Briefly, cells were lysed in 50 mM Tris-HCl pH 8.0, 150 mM NaCl, 5 mM EDTA, 1X complete protease inhibitor, and 1% n-dodecyl-β-D-maltoside, followed by centrifugation at 20,000 g for 1 hour at 4 °C. HLA-I–peptide complexes were immunoprecipitated from clarified lysates using anti-HLA-ABC antibody (W6/32, Bio X Cell) coupled to CNBr-activated Sepharose beads (GE Healthcare) with 18h incubation at 4 °C. After sequential washes, complexes were eluted with 0.25% TFA. Eluates were evaporated, resuspended in 30% ACN / 0.1% FA, and fractionated by strong cation exchange chromatography (PolyLC sulfoethyl A column) prior to LC-MS/MS analysis. Peptides were separated on a Dionex Ultimate 3000 system at 300 nL/min over a 0–30% ACN gradient (0.1% FA, 60 min) and analyzed on an Orbitrap Exploris 480 in data-dependent acquisition mode with HCD fragmentation, scanning 350–900 m/z.

LC-MS/MS spectra from HLA-I immunopeptidomics data^34^ were searched using FragPipe v23.1^105^ (MSFragger v4.3, Philosopher v5.1.2, Percolator v3.7.1) against the same custom FASTA database described above. Searches were performed with nonspecific cleavage, allowing peptide lengths of 7–12 amino acids and precursor masses of 200–5,000 Da. Precursor mass tolerance was ±20 ppm and precursor true tolerance was 15 ppm; fragment ion mass tolerance was 0.02 Da using b- and y-ion series, with a minimum of 5 matched fragments required per PSM. Intensity values were square root-transformed prior to scoring. Isotope errors of 0 and 1 were considered. Variable modifications included methionine oxidation (+15.995 Da), N-terminal protein acetylation (+42.011 Da), cysteine carbamidomethylation (+57.021 Da), N-terminal pyroglutamate from glutamine (−17.027 Da) and from glutamic acid (−18.011 Da), and cysteine propionamide (+71.037 Da). No fixed modifications were applied. PSM validation was performed using Percolator v3.7.1 with target-decoy competition (flags: --only-psms --no-terminate --post-processing-tdc). PSM-level FDR was controlled at 1% using group-specific FDR estimation in Philosopher, with PSMs grouped by protein existence (PE) level encoded in the FASTA headers — canonical UniProt entries (PE=1) and non-canonical translon-derived sequences (PE=2) were evaluated as separate groups, preventing FDR inflation from the larger canonical database. No protein-level FDR filter was applied. MSBooster v1.3.17 was used for DIA-NN-based retention time and spectral rescoring. Peptide sequences matching annotated canonical proteins were discarded. HLA binding prediction for class I alleles was performed using NetMHCpan 4.1^108^, with peptides classified as strong binders at a rank threshold of 0.5%.

LC-MS/MS spectra were analyzed by de novo peptide sequencing using Casanovo^109^ with the non-tryptic model weights. Spectra were processed retaining the 150 most intense peaks between m/z 50–2,500, with a minimum intensity of 0.01; precursor peaks within 2.0 Da were removed, and spectra with precursor charge >10 were excluded. Peptides were predicted with a minimum length of 8 residues and maximum length of 100 residues. Precursor mass tolerance was ±50 ppm, with isotope errors of 0 and +1 considered. Inference was performed on GPU with a batch size of 256 using greedy decoding (one beam) and one peptide prediction per spectrum. The residue vocabulary included carbamidomethylated cysteine, oxidized methionine, deamidated asparagine/glutamine, and the specified N-terminal modifications.

### Proteomics - synthetic peptide validation

Peptides were reconstituted in 0.015% n-Dodecyl-β-D-maltoside (DDM) before liquid chromatography-tandem mass spectrometry (LC-MS/MS) analysis. LC was performed on a Vanquish Neo LC system (Thermo Scientific) coupled to an Orbitrap Astral mass spectrometer, interfaced by a Nanospray Flex ion source (Thermo Scientific). Peptides were injected onto a Easy-SprayTM PepMapTM Neo UHPLC 150 μm x 15 cm or 50 cm low load- or 110 cm μPACTM HPLC columns (C18 column (75 µm inner diameter × 50 cm double nanoViper PepMap Neo, 2 μm, 100 Å), Thermo Scientific) regulated at 50 °C and separated with a linear gradient from 100% buffer A (100% H2O and 0.1% formic acid) to 28% buffer B (100% acetonitrile and 0.1% formic acid) at a flow rate of 300 nl min–1 over 60 min. The instrument was operated in Data-Dependent Acquisition (DDA). Peptides were then analyzed in MS applying a 2200V spray voltage, a funnel RF level of 40% and with a heated capillary temperature set to 285 °C. MS full scans were recorded using the Orbitrap mass analyzer in centroid mode for ranges 280-720 m/z with a resolution of 240,000, a normalized automatic gain control (AGC) target set at 300% and a maximum injection time (IT) of 100 ms.

### Random Forest classification and SHAP analysis

To assess the contribution of individual nc-translons to the separation of EwS from non-EwS samples, a Random Forest classifier was trained on the proteomics detection matrix from dataset 1 using scikit-learn 1.6.1. The classifier used 500 trees with balanced class weights to account for the imbalance between EwS and non-EwS samples. Model performance was evaluated using leave-one-out cross-validation (LOOCV), chosen given the small sample size. SHAP (SHapley Additive exPlanations) values were computed on the full-data model using the TreeExplainer implementation in the SHAP library 0.51.0, and taken for the EwS-positive class. Mean absolute SHAP values were used to rank features by their contribution to EwS classification.

### In silico peptide detectability analysis

To assess the reasons for differential detection of neogene translons by global proteomics versus immunopeptidomics, we performed an in silico peptide detectability analysis. For each neogene translon and a reference set of 364 random canonical UniProt proteins, we computed two metrics: tryptic peptide count and HLA strong binder count.

Tryptic digestion was performed in silico using a regular expression-based implementation of trypsin cleavage rules (cleavage after K or R, not before P), allowing up to 2 missed cleavages. Only peptides of 7–25 amino acids were considered detectable by mass spectrometry. The number of such peptides per protein was recorded as the tryptic peptide count.

HLA binding prediction was performed using NetMHCpan 4.1^108^. For each protein, all possible peptides of length 8–11 amino acids were generated by sliding window and submitted to NetMHCpan against HLA-A and HLA-B alleles present in the patient cohort. Peptides with an EL_Rank below 0.5 were classified as strong binders. The HLA strong binder count was defined as the total number of strong binder predictions summed across all alleles, where the same peptide binding to multiple alleles was counted multiple times.

### poly(A) readthrough analysis

To identify ribosome-protected fragments (RPFs) spanning the exon–poly(A) junction, we constructed short reference sequences consisting of the last 30 nucleotides of each MANE Select protein-coding transcript followed by 30 As, representing the poly(A) tail. These sequences were indexed using bowtie2^87^ (v2.4.2) with default parameters. Trimmed Ribo-seq reads were aligned to this index using bowtie2^87^ with a seed length of 25 (*`--seedlen=25`*), retaining both mapped (*`--al-gz`*) and unmapped (*`--un-gz`*) reads. Only uniquely mapping reads (MAPQ = 255) that spanned the exon–poly(A) junction were considered for downstream analysis.

To distinguish true poly(A) readthrough events from reads mapping to A-rich exonic sequences, we applied three filters. First, we retained only reads with at least five terminal adenosine. Second, we verified the transcript-specificity of each read by stripping trailing adenosines from the read sequence and aligning the remaining exonic portion against the full Ensembl (v113) transcriptome using BLASTn (v2.14.0; *`-word_size 7 -evalue 1000`*). Only reads whose exonic portion produced a perfect full-length match to a single gene were retained as high-confidence poly(A) readthrough events. Lastly, we disregarded genes that had less than 25% of the reads to be uniquely mapping, to only retain genes with a significant amount of poly(A) mapping.

### Transfection of transient overexpression vectors of neogene and neoisoform translons

To transfect MHH-ES-1/TR/shEF1 cells with transient overexpression vectors containing neogene or neotranscript translons, 660,000 cells per condition were harvested, washed twice with PBS and dissociated using TrypLE (Gibco). Cells were centrifuged at 90 g for 10 min at room temperature and resuspended in SG nucleofection solution (Lonza), prepared by mixing 16.4 µL nucleofector solution with 3.6 µL supplement per reaction. 1.32 µg of pEF1α_T7_custom plasmid (GeneWiz, **Figure S11a**) encoding each FLAG-tagged translon was combined with 22 µL of cell suspension in a 96-well Nucleocuvette strip and nucleofected using program EH-100 on the 4D-Nucleofector (Lonza). As a positive control for the transfection and downstream experiments, a plasmid containing EGFP-FLAG was also taken along. After nucleofection, 80 µL of pre-warmed complete culture medium was added and the strip was incubated at 37 °C for 10 min before transferring cells to 12-well plates.

### Transient neoprotein detection

To measure FLAG+ cells, we used flow cytometry of MHH-ES-1/TR/shEF1 cells nucleofected as described above with 0.66 µg of plasmid per condition. 48h after nucleofection, cells were harvested, washed with PBS and transferred to a V-bottom 96-well plate. Cells were stained with eFluor 780 fixable viability dye (eBioscience) diluted 1:1000 in PBS for 10 min at 4 °C protected from light, then washed with FACS buffer. Cells were fixed and permeabilized using the eBioscience Foxp3/Transcription Factor Staining Buffer Set (cat# 00-5523-00) according to the manufacturer’s instructions: cells were incubated in 50 µL of 1X Fixation/Permeabilization solution for 30 min at 4 °C, then washed with 1X Permeabilization Buffer. Intracellular FLAG-tag detection was performed by incubating cells with CoraLite Plus 647-conjugated anti-FLAG monoclonal antibody (Proteintech) diluted in 1X Permeabilization Buffer for 30 min at 4 °C. After washing with Permeabilization Buffer and FACS buffer, cells were resuspended in 150 µL FACS buffer and acquired on a BD FACSymphony A1 flow cytometer (BD Biosciences). Compensation was calculated using single-stained controls for GFP (488 nm laser, 530/30 filter), CoraLite 647 (640 nm laser, 670/30 filter) and eFluor 780 viability dye (640 nm laser, 780/60 filter).

### Endogenous tagging of neoproteins

To enable CRISPR-Cas9-mediated homology-directed repair (HDR) for knock-in (KI) of a FLAG-P2A-mNeonGreen reporter cassette, a double-stranded donor DNA template was synthesized (Twist Bioscience, CA, USA). The donor construct was designed to insert the reporter in frame at the target locus and consisted of a left homology arm (LHA), a GSG linker, the FLAG-P2A-mNeonGreen cassette, and a right homology arm (RHA). The synthesized DNA was assembled into linearized pJET2.1 vector (Thermo Scientific™, MA, USA) using the HiFi DNA Assembly Cloning Kit (New England Biolabs, MA, USA). Correct assembly was validated by Sanger-sequencing using pJET1.2 reverse sequencing primer (5′-AAGAACATCGATTTTCCATGGCAG-3′).

Universal trans-activating CRISPR RNA (tracrRNA, 67mer, 5 µL) (Integrated DNA Technologies, IA, USA), dissolved in nuclease-free duplex buffer (100 µmol/L) was added to custom-designed synthesized CRISPR RNA (crRNA, 5 µL) (Integrated DNA Technologies) encoding a binding sequence specific to the target gene (5′-AGGGAACGAGCTAATCAATT-3′) dissolved in IDTE (pH 7.5, 100 µmol/L). For guide RNA (gRNA) formation tracrRNA and crRNA were duplexed in a thermocycler by initial denaturation at 95 °C for 5 min, followed by 90 cycles of 12 s each, beginning at 98 °C and decreasing by 1 °C per cycle. Next, ribonucleoprotein (RNP) composed of active CRISPR associated protein 9 (Cas9, 62 µmol/L) (Integrated DNA Technologies) and gRNA duplex (50 µmol/L) was generated (6 µL, final concentration of 20 µmol/L) and incubated at room temperature (20 min). The resulting RNP complex was then used for one electroporation reaction to induce CRISPR-Cas9 mediated HDR knock-in. Next, A-673 wild-type cells (1×10□) were resuspended in electroporation buffer (18 µL) composed of SF Cell Line Nucleofector® Solution (14 µL) and Supplement 1 (4 µL) (Lonza Group AG, Switzerland). RNP (6 µL) and DNA donor template (2 µg) were added to the cells, which were electroporated (CM-130 pulse code, SF buffer) in 16-well strips using the 4D-Nucleofector (Lonza Group AG). After electroporation cells were incubated at room temperature (10 min) and then transferred to prewarmed complete cell culture medium (20% FCS). Following electroporation (12 h) cell culture medium was replaced with fresh complete medium (20% FCS). 14 days after electroporation cells expressing mNeonGreen (green) were sorted into 96-well plates containing complete cell culture medium (20% FCS) using FACSAria Fusion cytometer (Becton Dickinson, NJ, USA). Wells were monitored every third day under an inverted phase-contrast microscope to confirm the presence of single-cells. Wells containing multiple cells were excluded from further analysis. Clonal outgrowth was monitored over the course of three weeks. Once a cell clone reached 90% confluence, cells were carefully transferred to 24-well plates for expansion. Correct KI was validated by PCR amplification of the edited genomic locus from genomic DNA of single-cell clones using forward primer (5′-CAGGGTTGCTGTGGATTACA-3′) and reverse primer (5′-GGTCTCCAGAAACTTAGGAAAGG-3′) flanking the Cas9 cleave site and subsequent Sanger-sequencing. Sequence validated clones were expanded for functional assays.

### Subcellular localization staining

For immunofluorescence, MHH-ES-1/TR/shEF1 cells were nucleofected as described above with 0.66 µg of plasmid per condition. After nucleofection, cells were seeded into µ-Slide 8-well chambers (ibidi) at approximately 10,000 cells per chamber in 30 µL and allowed to adhere for 30 min at 37 °C before adding 100 µL of complete culture medium. Cells were cultured for 48h before staining.

Cells were washed with ER buffer and permeabilized with 0.1% Triton X-100 in ER buffer for 1 min at room temperature. After washing, non-specific binding was blocked using Image-iT FX Signal Enhancer (Invitrogen) for 30 min at room temperature, followed by 5% normal goat serum in PBS for 1h at room temperature. Two antibody panels were used: Panel 1 included anti-FLAG (mouse) and anti-ATPIF1 (rabbit, mitochondria); Panel 2 included anti-FLAG (mouse) and anti-Golgin-97 (rabbit). Primary antibodies were diluted in 1% BSA/PBS and incubated overnight at 4 °C. After washing, cells were incubated with Alexa Fluor 555-conjugated goat anti-mouse and Alexa Fluor 750-conjugated goat anti-rabbit secondary antibodies diluted in 0.2% BSA/PBS for 1h at room temperature protected from light. Cells were then washed and stained with 1X ER-Tracker Red (Invitrogen) for 1h at 37 °C in the dark. For Panel 1, cells were subsequently stained with 1X Golgi-Tracker Green for 1h at 4 °C in the dark, followed by a re-incubation step in complete DMEM for 45 min at room temperature. Slides were mounted using SlowFade Diamond Antifade Mountant with DAPI (Invitrogen) and imaged on a Stellaris confocal microscope (Leica).

A-673 wild-type cells and A-673 cells carrying an endogenous FLAG-P2A-mNeonGreen knock-in at the Ew_NG157 locus were seeded in chamber slides (ibidi) and cultured for 24 h. Cells were washed with PBS and fixed with freshly prepared 4% formaldehyde in PBS for 15 min at room temperature. Following fixation, cells were washed with cold PBS and permeabilized with 0.1% Triton X-100 in PBS for 10 min at room temperature. Samples were washed again with cold PBS and blocked with 1% BSA in PBS for 1 h at room temperature. For detection of FLAG-tagged protein, samples were incubated overnight at 4 °C with monoclonal anti-FLAG antibody (Proteintech, Clone 8H6A10) diluted 1:200 in antibody dilution buffer containing 1% BSA and Tween-20 in PBS. The following day, samples were washed with cold PBS and incubated with the appropriate fluorophore-conjugated secondary goat anti-mouse Alexa Fluor 647 antibody for 2 h at room temperature protected from light. After washing, nuclei were counterstained with DAPI, and slides were mounted using Fluoroshield mounting medium. Samples were stored at 4 °C protected from light until imaging.

### Cell growth assay

After nucleofection with 0.66 µg of transient overexpression plasmid as described above, approximately 330,000 cells per condition were seeded in triplicates in a 48-well plate (Greiner Bio-One) containing 370 µL of pre-warmed complete culture medium. Cell growth was then measured by taking HD Phase images every 6 h at 4 positions per well using the Incucyte (Sartorius), calculating average confluence for each timepoint over 4 d.

### siRNA knockdown of neogenes

EwS A-673 and MHH-ES1 cells were reverse-transfected with siPOOLs (siTOOLS Biotech; siRNA sequences listed in **Supplementary Table S11)** using Lipofectamine RNAiMAX (Thermo Fisher) according to the manufacturer’s instructions with minor modifications. Transfections were performed in 6-well plates at a final volume of 2 mL complete growth medium (RPMI, 10%FCS, 1%P/S) per well. For each well, 40 µL siPOOL stock solution (150 nM) was diluted in 246 µL Opti-MEM (Thermo Fisher) together with 4 µL RNAiMAX reagent to generate a total transfection mixture volume of 500 µL per well. The transfection mixture was gently mixed and incubated for 10–20 min at room temperature to allow formation of lipid–RNA complexes. Cells were prepared as a single-cell suspension at a density of 2 × 105 cells/mL, and 300,000 cells in 1.5 mL complete growth medium (RPMI, 10%FCS, 1%P/S) were added to each well containing the transfection complexes, resulting in a final volume of 2 mL per well. Cells were maintained under standard culture conditions (37 °C, 5% CO□) for 72 h prior to EdU-based proliferation analysis.

For EdU incorporation assays, cells were incubated with 10 µM EdU for 4-8 h prior to harvest. Cells were washed once with PBS, detached using trypsin, and fixed in 4% paraformaldehyde (PFA) for 15 min at room temperature. Fixed cells were washed with 1% BSA in PBS and permeabilized with 0.1% (v/v) Triton X-100 in PBS for 30 min. Click-iT^TM^ detection was performed according to the manufacturer’s instructions using either Alexa Fluor 488 azide or Alexa Fluor 647 picolyl azide. Alexa Fluor 647 picolyl azide was used when Click-iT labeling was combined with propidium iodide (PI)-based cell cycle analysis. Flow cytometric measurements were acquired on a BD FACSCanto II (BD Biosciences), and acquisition was stopped after recording 10,000 single cells per sample. Data were analyzed using FlowJo software (version 10; FlowJo LLC).

### Knockdown shRNA against Ew_NG48

Lentiviral particles carrying the shRNAs targeting Ew_NG48 (sh17985, sh17986 and sh17987, **Supplementary Table S11**) or a non-targeting control shRNA (SHC002, Merck-Sigma) were produced in HEK293T cells. Briefly, 10^7^ HEK293T cells were seeded in T75 flasks one day prior to transfection. Cells were co-transfected with the vector carrying the shRNA of interest, the packaging plasmid pPAX2 and the envelope plasmid VSV-G using Lipofectamine 2000 (Thermo Fisher Scientific). For each transfection, 10 μg of pPAX2, VSV-G and shRNA plasmids were diluted in Opti-MEM and combined with Lipofectamine 2000 according to the manufacturer’s instructions. After a 15-min incubation at room temperature, transfection complexes were added dropwise to HEK293T cells cultured in antibiotic-free medium. The culture medium was replaced the following day with complete DMEM supplemented with 10% FBS, 1% penicillin/streptomycin and GlutaMAX. Viral supernatants were collected 48h post-transfection and filtered through 0.45-μm filters and titrated by transducing 200,000 A673/TR/shEF1 cells with serial dilutions of lentiviral supernatants (10^-1^ to 10^-5^). A673/TR/shEF1 cells were subsequently transduced with each lentiviral preparation at multiplicities of infection (MOIs) of 1 and 5. GFP-positive cells were isolated by fluorescence-activated cell sorting (FACS). A673/TR/shEF1 cells transduced at MOI 1 were selected for further experiments with all shRNA constructs (sh17985, sh17986 and sh17987) to minimize the risk of multiple vector integrations while maintaining efficient shRNA induction. A673/TR/shEF1 cells were either treated with doxycycline to induce shRNA targeting EF1 or with 0.5 mM IPTG to induce shRNA against Ew_NG48 for 3 or 6 days. Ew_NG48 and EF1 knockdown efficiency was assessed by RT-qPCR.

### CRISPRi targeting Ew_NG48

A-673 expressing dCas9-KRAB^19^ (A673-dCas9-KRAB) was used to assess CRISPRi inhibition against Ew_NG48. Regions flanking Ew_NG48 FLI1-bound GGAA microsatellite promoter or enhancer regions were targeted using guides (**Supplementary Table S12**). To this aim, 50’000 A-673-dCas9-KRAB cells were seeded in 6-well plates. The following day, A-673-dCas9-KRAB cells were incubated with a mix of OptiMEM (Gibco), RPMI 1640 medium (Sigma) supplemented with 10% FBS (Eurobio), RNA guides and track RNA (IDT DNA) at a final concentration of 10nM for 4 days. After 4 days, RNA was extracted using a NucleoSpin RNA kit (Macherey-Nagel) and quantified using a Nanodrop device. Ew_NG48 expression quantification was assessed using RT-qPCR as described below.

### RT-qPCR

Reverse transcription was achieved using Applied Biosystems High-Capacity cDNA Reverse Transcription kit with RNase Inhibitor (Fisher Scientific) according to the manufacturer’s instructions. qPCR was performed in triplicates using SYBR Green PCR Master Mix (Fisher Scientific) and the CFX384 Touch Real-Time PCR System and analyzed with CFX Manager Software. *RPLP0* was used as a reference housekeeping gene for normalization.

## Acknowledgements

We acknowledge the Utrecht Sequencing Facility (USEQ) for providing sequencing service and data. USEQ is subsidized by the University Medical Center Utrecht and The Netherlands X-omics Initiative (NWO project 184.034.019). We thank Alex Kentsis, Eliyahu Havasov, Katarzyna Kulej, and Asher Preska Steinberg for providing early-access to and expertise on the analysis of their A-673 multiprotease proteomics data. High-throughput sequencing was also performed by the ICGex NGS platform of the Institut Curie supported by the grants ANR-10-EQPX-03 (Equipex) and ANR-10-INBS-09-08 (France Génomique Consortium) from the Agence Nationale de la Recherche (“Investissements d’Avenir” program), by the ITMO-Cancer Aviesan (Plan Cancer III) and by the SiRIC-Curie program (SiRIC Grant INCa-DGOS-465 and INCa-DGOS-Inserm_12554). Data management, quality control and primary analysis were performed by the Bioinformatics platform of the Institut Curie. We would like to express our sincere thanks to Carsten Maus, Erjia Wang (Genomics and Proteomics Core Facility, DKFZ). Lena Weiser, Gregor Warsow (Omics IT and Data Management Core Facility, DKFZ) for their highly dedicated support in data management and processing. Robert Autry, Gnanaprakash Balasubramanian, Christopher Previti and Rolf Kabbe (Division of Pediatric Neurooncology, DKFZ) for their sincere and dedicated contribution to the bioinformatics analyses.

This project was partially supported by the Fight Kids Cancer Funding Programme (“NEWtargets – Recurrent EwS-specific Neoproteins as Targets for Immunotherapy”), supported by Imagine for Margo, Fondation KickCancer, Fondatioun Kriibskrank Kanner, Federazione Italiana Associazioni Genitori e Guariti Oncoematologia Pediatrica, Cris Cancer Foundation and Stichting Kinderen Kankervrij (KiKa), to S.v.H., T.G.P.G., O.D., and J.J.W. S.v.H. acknowledges funding from Stichting Reggeborgh. Research reported in this publication was supported by Oncode Accelerator, a Dutch National Growth Fund project under grant number NGFOP2201, awarded to S.v.H. This work is co-financed by Oncode Institute, which is partly funded by the Dutch Cancer Society. The further development and translation of this project will be supported by the ILLUMINE team through the Cancer Grand Challenges partnership funded by Cancer Research UK (CGCATF-2025/100034), the National Cancer Institute, the Cancer Research Institute, and KiKa (Children Cancer Free Foundation) and the Cancer Grand Challenges KOODAC^3^ grant n°CGCATF-2023/100017 funded by Cancer research UK, KiKa (Children Cancer Free Foundation) and INCa. This work was supported by the European Union’s Horizon Europe research and innovation programme through the Marie Skłodowska-Curie Actions Staff Exchanges (MSCA-SE) Grant Agreement No. 101182922 (GRASSHOPPER). J.L. acknowledges support by the German Cancer Aid (DKH-70115914). T.G.P.G. acknowledges funding by the Dr. Leopold und Carmen Ellinger foundation, the Ministry of Education and Research (BMBF; SMART-CARE), and the European Union (ERC, CANCER-HARAKIRI, 101122595). Views and opinions expressed are however those of the authors only and do not necessarily reflect those of the European Union or the European Research Council (ERC). The INFORM program is financially supported by the German Cancer Research Center (DKFZ), several German health insurance companies, the German Cancer Consortium (DKTK), the German Federal Ministry of Education and Research (BMBF), the German Federal Ministry of Health (BMG), the Ministry of Science, Research and the Arts of the State of Baden-Württemberg (MWK BW); the German Cancer Aid (DKH), the German Childhood Cancer Foundation (DKS), RTL television, the aid organization BILD hilft e.V. (Ein Herz für Kinder) and the generous private donation of the Scheu family. W.F. acknowledges support from SiRIC-3: INCa-DGOS-Inserm-ITMO cancer_18000. U.D. acknowledges support from the German Cancer AID (70113419) and German Federals Minitry of Eductation and Research (HEROES AYA 01KD2207B).

F.H.G. acknowledges support by the KiTZ Foundation in memory of Kirstin Diehl and the Henrik-Kreibohm-Foundation, and by the German Cancer Aid through the ‘Mildred-Scheel-Doctoral Program and the German Academic Scholarship Foundation. J.J.W. acknowledges support from the Fight Kids Cancer and St Baldrick’s Foundation Arceci Innovation Award. We are very grateful to the following associations for providing essential support: L’Etoile de Martin, la Course de l’Espoir, M la vie avec Lisa, ADAM, Couleur Jade, Dans les pas de Géant, Courir pour Mathieu, Marabout de Ficelle, Olivier Chape, Les Bagouzamanon, Enfants et Santé, Les Amis de Claire, Un Elan pour Lucas, and Amarape. O.D. acknowledges funding from Ligue contre le cancer grant AAEAC2020.LCC/OD, and Institut National du Cancer (INCa) high risk high gain grant (convention N°2023-022). M.K.K. was supported by a fellowship from the Eureka Foundation. S.v.H., T.G.P.G. and J.H.M.M. acknowledge support by EU-CAN-KIDS that stimulates collaborations and joint projects between the Hopp Children’s Cancer Center (KiTZ) in Heidelberg, the Princess Máxima Center for Pediatric Oncology in Utrecht and the Institut Curie/lnserm in Paris.

## Author Contributions

S.v.H., O.D., T.G.P.G., J.J.W., L.A.B., J.P., M.K.K.: conceptualization, methodology, writing - original draft, review & editing. S.v.H., O.D., T.G.P.G., J.J.W.: funding acquisition, supervision. L.A.B., J.P., and M.K.K.: validation, investigation, formal analysis, visualization. J.L., F.H.G., M.J.C.-G., N.H., N.G., S.G., E.G.A.W., O.J., L.S., T.L., S.A.G.E., E.N., A.P.L., M.V., W.F., A.I.L., K.L.-D., J.M.M., C.M., C.H.: investigation. F.A.: formal analysis, conceptualization, supervision. J.K., J.H.M.M., A.v.O.: supervision, resources. M.I.S., H.T., E.V., G.W.O., B.L., U.D.: resources. I.J., S.B., S.L.: methodology, investigation. J.L., F.H.G., S.G., E.G.A.W., K.L.-D.: methodology. L.S., K.A.: formal analysis. O.L. investigation, review & editing.

## Declaration of generative AI and AI-assisted technologies

During the preparation of this work, the authors used commercial LLMs to assist with the writing of code and manuscript text. After using this tool or service, the authors reviewed and edited the content as needed and take full responsibility for the content of the publication.

## Supplementary Tables

**Supplementary table S1**. Genomic coordinates in hg38 of all 124 neogenes in gtf format.

**Supplementary table S2**. Neogenes annotation, characteristics and fold-change expression in doxycycline-inducible cellular models.

**Supplementary table S3**. Genomic coordinates in hg38 of all 2,244 neoisoforms and their translons in gtf format.

**Supplementary table S4**. Cohort description of the 48 EwS samples analyzed by Ribo-seq.

**Supplementary table S5**. Overview of all non-canonical translons identified in Ribo-seq data from EwS patient samples, including their characteristics and associated statistics.

**Supplementary table S6**. Overview of 2,244 neoisoforms identified with long-read RNA-seq with their characteristics and associated translons annotation.

**Supplementary table S7**. Neogene- and neoisoform-derived translons and non-canonical ORFs detection in global proteomics and immunopeptidomics datasets.

**Supplementary table S8**. Neogene- and neoisoform-derived translons and non-canonical ORF peptides detected in global proteomics and immunopeptidomics datasets.

**Supplementary table S9**. Overview of evolutionary characteristics of neogene and neoisoform translons and non-translated control ORF sequences (NT controls).

**Supplementary table S10**. Overview of neogene and neoisoform translons selected for overexpression experiments.

**Supplementary table S11**. Complete list of all siRNA sequences included in siPOOLS.

**Supplementary table S12**. Primers, guides and probes sequences used for shRNA, CRISPRi and qPCR experiments.

## Supplementary Figures

**Figure S1.**
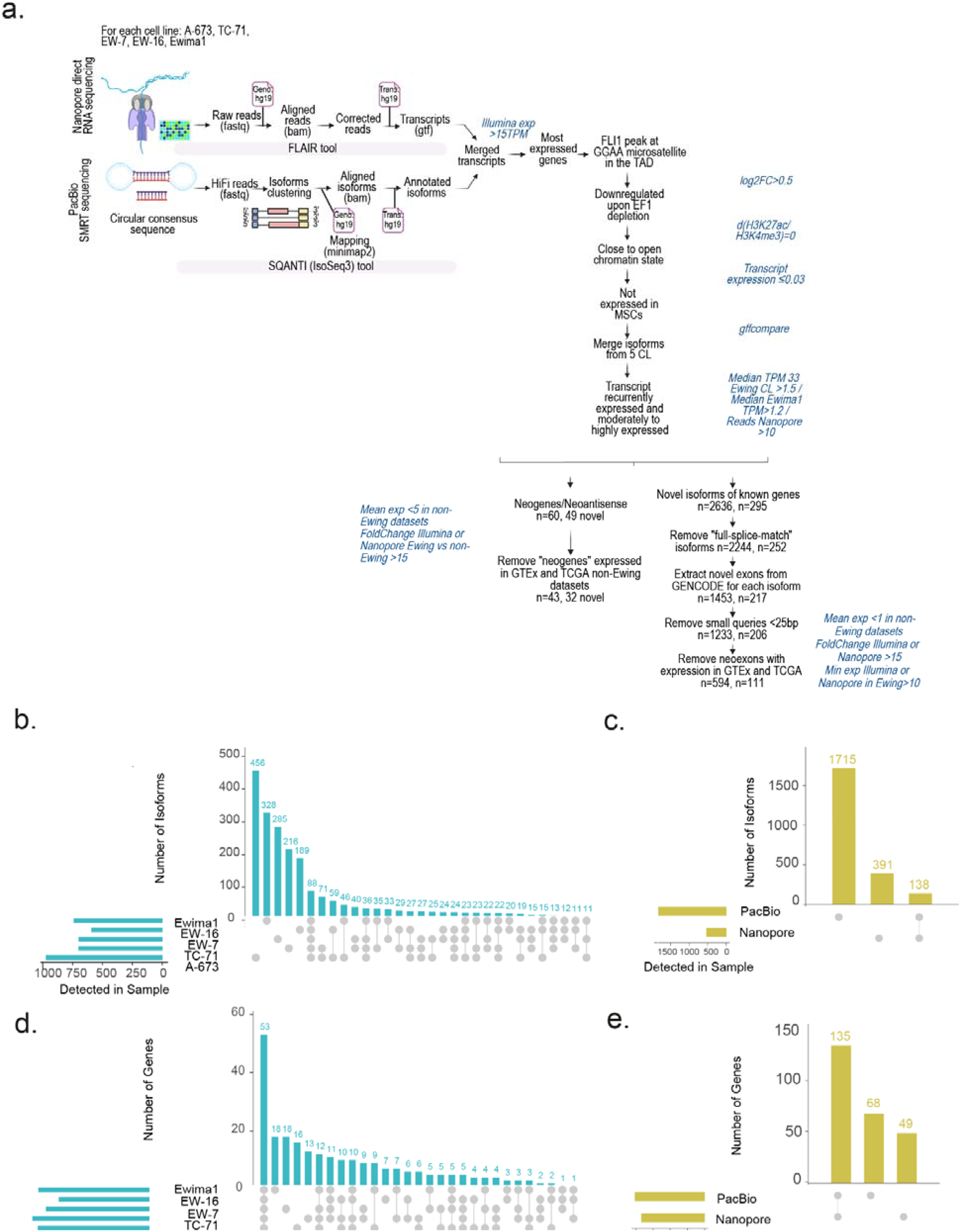
Pipeline and detected genes and isoforms in long-read RNA-seq technologies. **a**. Overview of the long-read RNA-seq pipeline for neoisoforms and neogenes discovery. MSCs: Mesenchymal stem cells; CL: cell lines. **b-e**. Number of isoforms (**b**) and genes (**d**) detected in the 5 EwS cell lines models. Number of neoisoforms (**c**) and genes (**e**) detected in each long-read sequencing technology PacBio and Nanopore.

**Figure S2.**
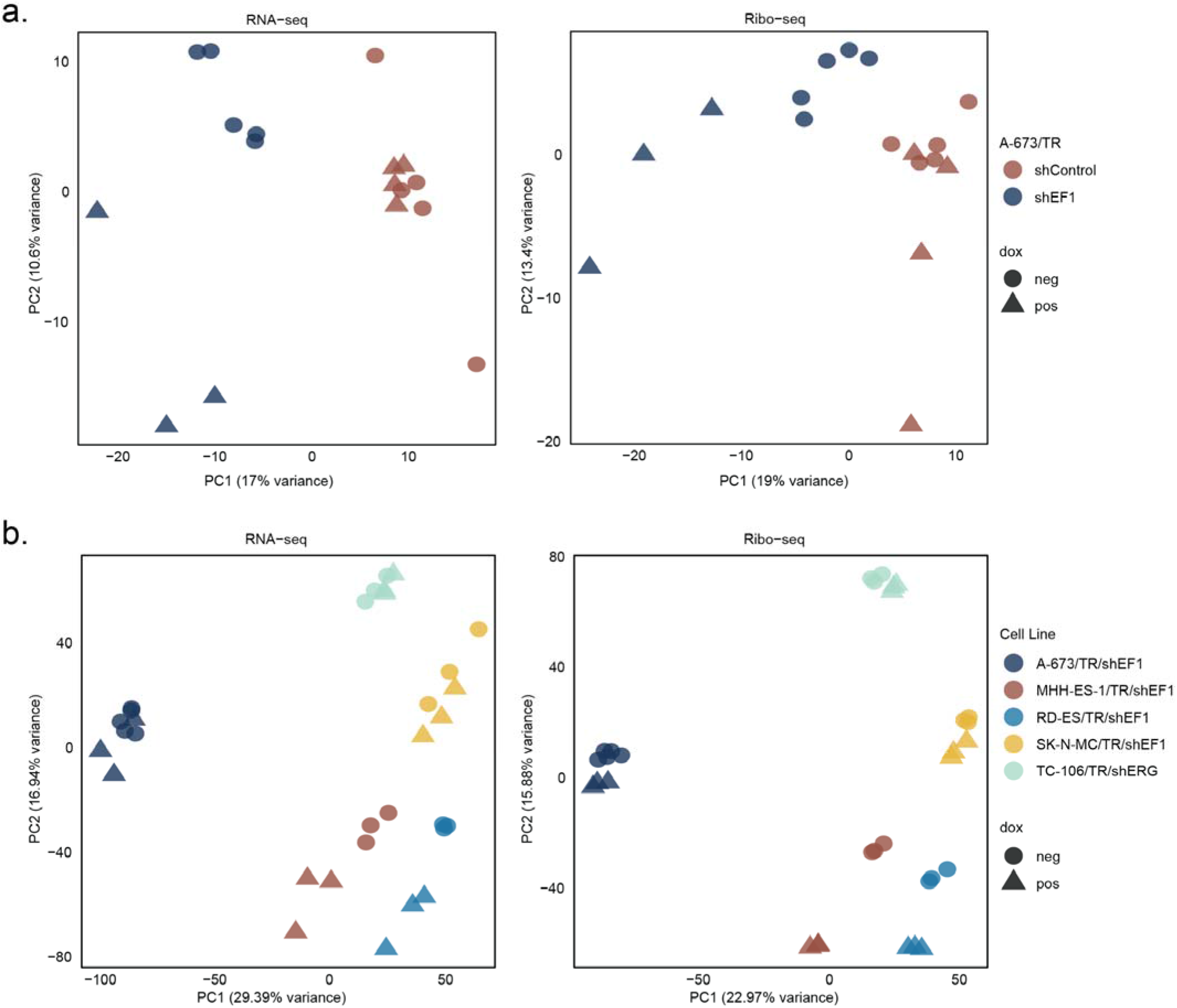
Principal component analysis (PCA) plots for EwS dox-inducible cell lines. **a.** PCA plots on th RNA-seq and Ribo-seq level comparing the effect of dox-induction in A-673/TR/shEF1 vs A-673/TR/shControl. **b.** PCA plots on the RNA-seq and Ribo-seq level for the different dox-inducible EwS cell lines.

**Figure S3.**
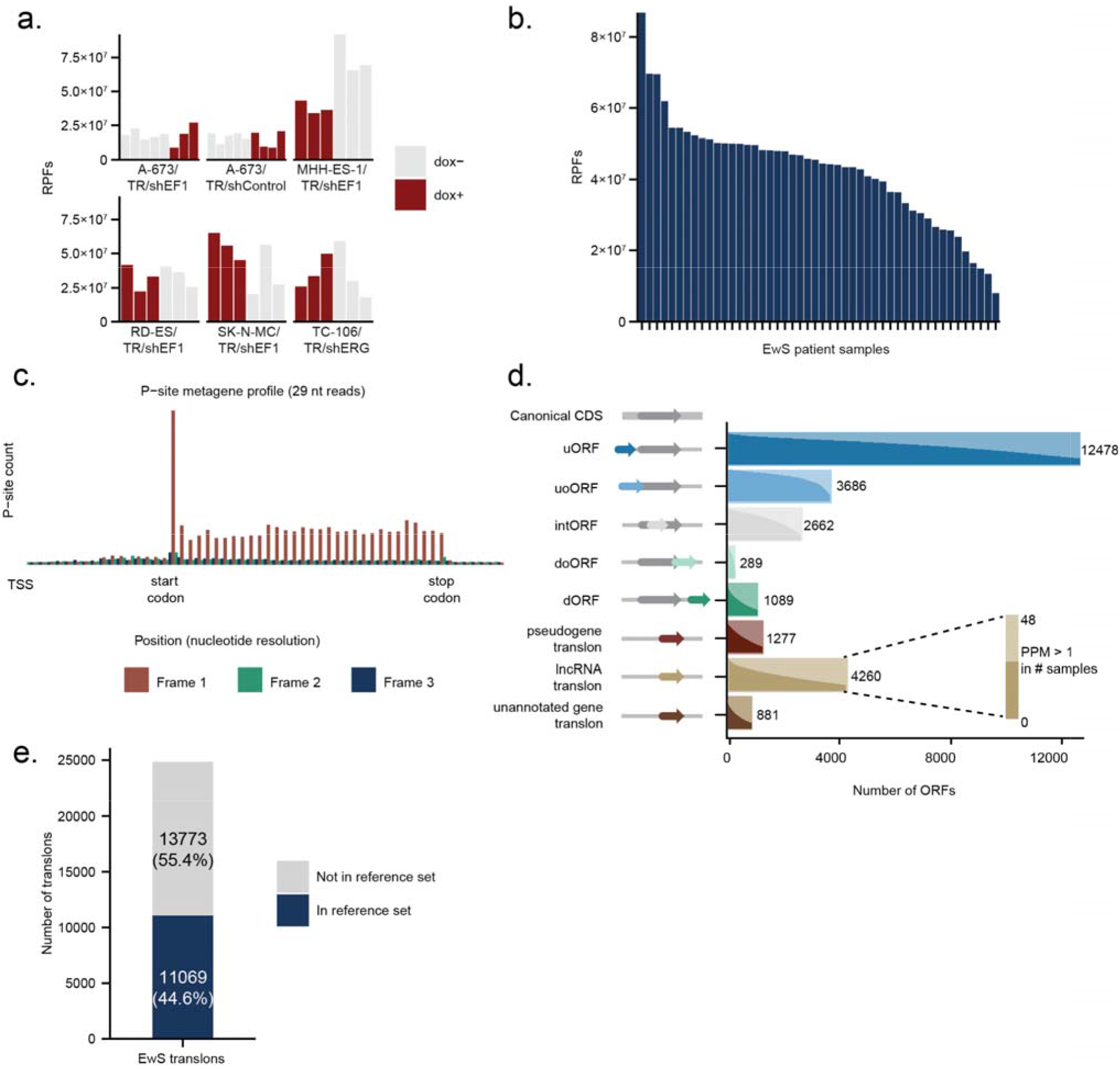
Ribo-seq QC statistics and all non-canonical translon calls. **a.** Barplot indicating the number of ribosome protected fragments (RPFs) after mapping for each of the EwS cell line samples. **b.** Barplot indicating the number of RPFs after mapping for each of the 48 EwS patient samples. **c.** Metagene plot for 29nt reads in the EwS patient Ribo-seq samples. **d.** Overview of the different nc-translon biotypes called in our EwS dataset (see Methods for details). The accompanying barplot shows the number for each of the nc-translon biotypes, with the histogram showing the recurrence of translation for each translon. **e.** Barplot showing the overlap of non-canonical translons found in our EwS dataset vs the transCODE human nc-translon reference catalog^42^.

**Figure S4.**
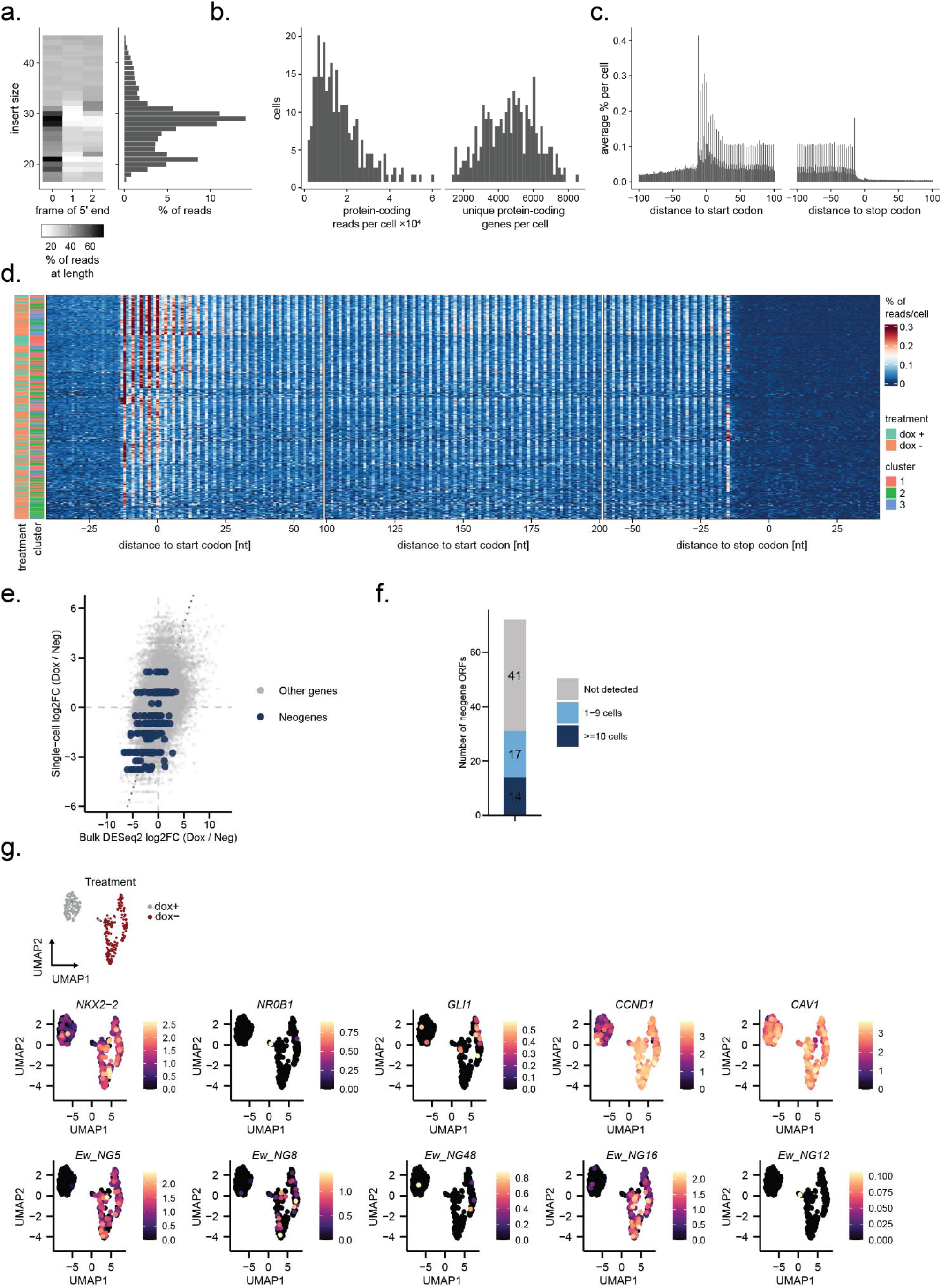
Supplementary information for single-cell Ribo-seq (scRibo-seq) **a.** Heatmap and histogram showing read length distribution. **b.** Histograms showing the distribution of the total number of reads (cell_total) and genes (num_genes) per cell. **c.** Metagene plot showing the frame preference for all combined scRibo-seq reads. **d.** Heatmap showing the frame preference across cells and conditions. **e.** Scatterplot depicting the correlation between the translation levels after doxycycline induction in scRibo-seq and bulk Ribo-seq of MHH-ES-1/TR/shEF1 cells. **f.** Barplot showing the number of neogenes for which we found reads in the scRibo-seq data **g.** UMAP projection showing the scRibo-seq expression of 5 highly translated neogenes and 5 well-established fusion targets.

**Figure S5.**
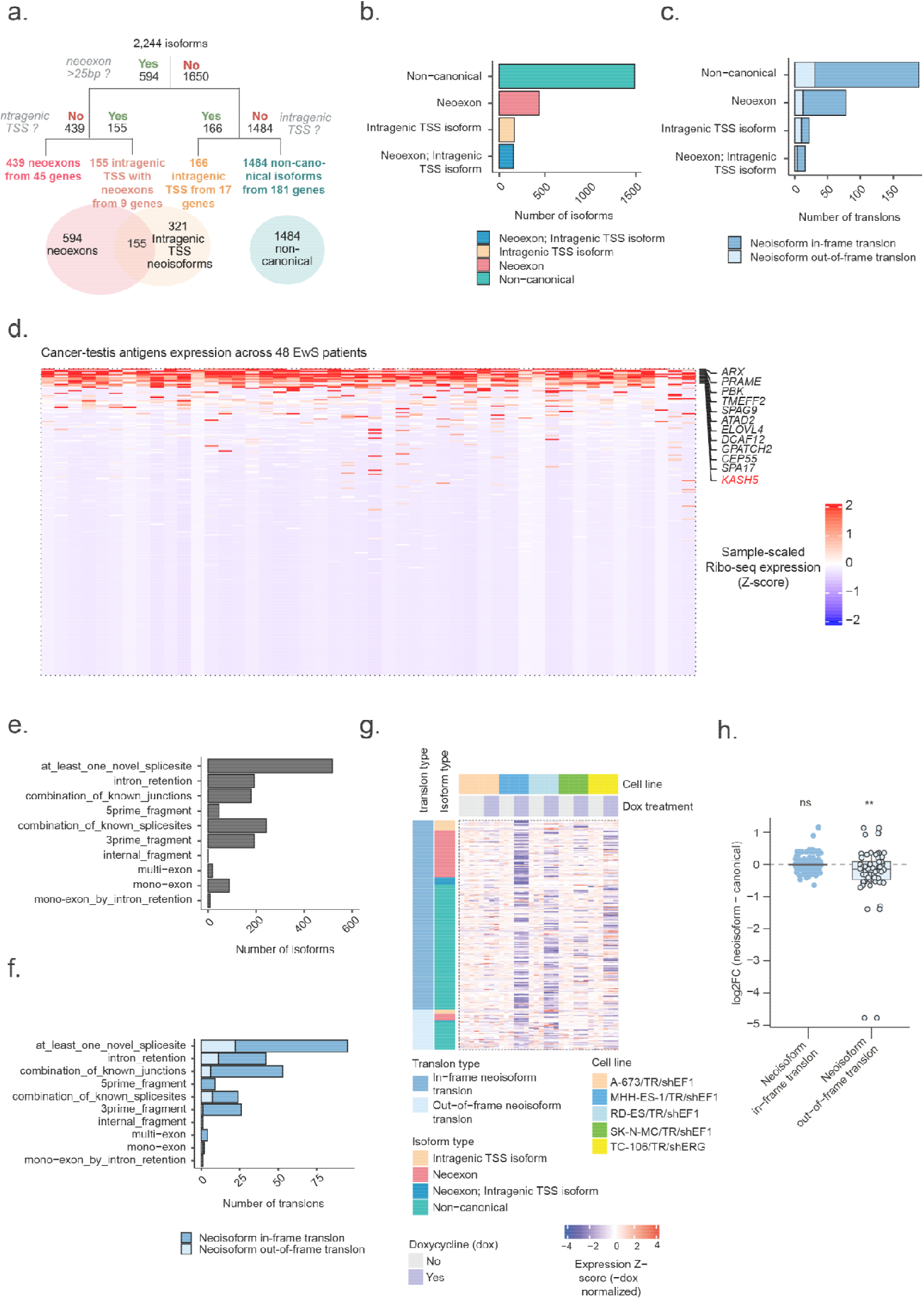
EwS-specific neoisoforms and associated translons description. **a.** Schematics representing EwS-specific neoisoforms classification according to (i) Neoexon inclusion, (ii) Intragenic TSS and (iii) other splicing events. **b-c.** Number of isoforms (**b**) and associated translons (**c**) for each of the EwS-specific neoisoforms categories. **d.** Heatmap showing Ribo-seq expression Z-score of 239 annotated cancer testis antigens across 48 patients EwS tumors. *KASH5* is among the most expressed CTAs in EwS patient tumors. **e.** Unsupervised annotation (Sqanti) of splicing events occurring in 1484 EwS-specific non-canonical neoisoforms. **f**. Number of translons produced by each of the isoform categories for 1484 EwS-specific non-canonical neoisoforms. **g.** Heatmap showing Riboseq expression Z-score normalized with -dox condition in 5 EwS cell lines harboring shRNA against EWSR1::FLI1 or EWSR1::ERG, respectively. Row annotation shows translons and isoform subtypes. Column annotation highlights EwS cell lines and doxycycline treatment. There are 5 replicates for the A-673/TR/shEF1 -dox condition (far left samples), and 3 replicates for all other conditions. **h**. Boxplot showing Ribo-seq fold-change in expression of neoisoform translons (in-frame and out-of-frame) relative to canonical isoforms following knockdown using dox-inducible shRNA targeting *EWSR1::FLI1* or *EWSR1::ERG*, respectively, across five EwS cell lines. ** indicate a p < 0.01 (Wilcoxon signed-rank test).

**Figure S6.**
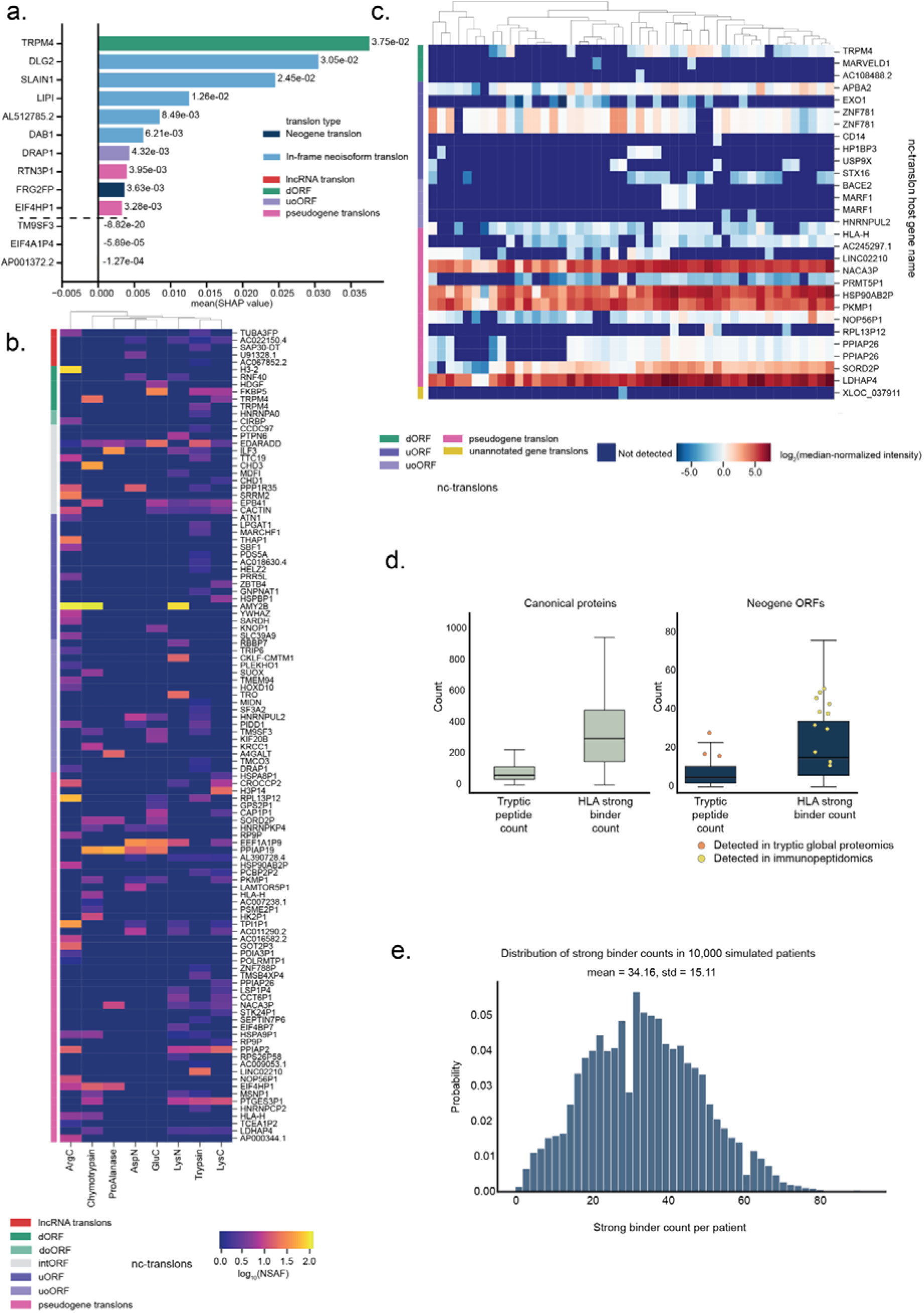
Proteomics analyses of neogene, neoisoform, and nc-translons. **a.** Mean SHAP values from a Random Forest classifier trained on the data from Figure 4c, showing the top 10 features by absolute value. Positive values indicate features driving EwS classification; negative values indicate features associated with non-EwS controls. Features are colored by translon type. **b.** Heatmap of nc-translons (excluding neogenes and neoisoforms) detected in the A-673 deep multienzyme digestion dataset^33^. Values represent log-transformed NSAF. **c.** Heatmap of nc-translons (excluding neogenes and neoisoforms) detected across 47 EwS patient tumor samples. Values represent log-transformed NSAF. **d.** Comparison of predicted tryptic peptide counts and HLA strong binder counts for canonical proteins (left) and neogene translons (right). Individual neogene translons detected in tryptic global proteomics or immunopeptidomics are overlaid as colored dots. The higher ratio of HLA strong binders to tryptic peptides in neogene translons compared to canonical proteins provides a mechanistic explanation for their enrichment in immunopeptidomics data. **e.** Predicted HLA coverage of the neogene and neoisoform translon set across a simulated patient population. HLA allele frequencies representative of the French population were used to simulate 10,000 patients. The distribution of predicted strong binders per patient is shown, illustrating the potential breadth of population coverage of the identified translon set as vaccine targets.

**Figure S7.**
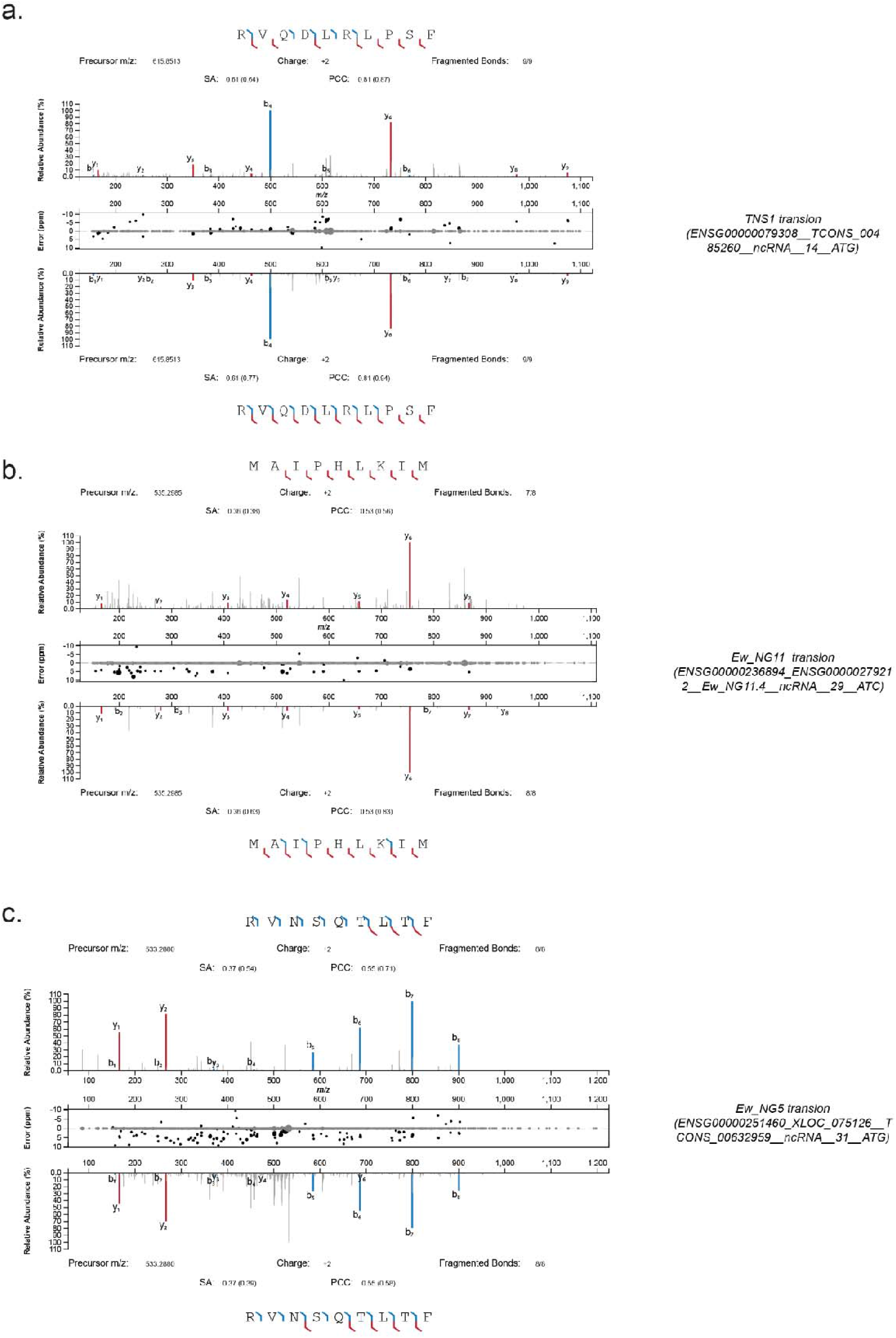
Synthetic spectra validation of the HLA-presented neogene and neoisoform peptides. **a.** *TNS1* peptide derived from translon ENSG00000079308 TCONS_00485260 ncRNA 14 ATG, **b.** Ew_NG11 peptide derived from translon ENSG00000236894_ENSG00000279212 Ew_NG11.4 ncRNA 29 ATC and **c.** Ew_NG5 peptide derived from translon ENSG00000251460_XLOC_075126 TCONS_00632959 ncRNA 31 ATG.

**Figure S8.**
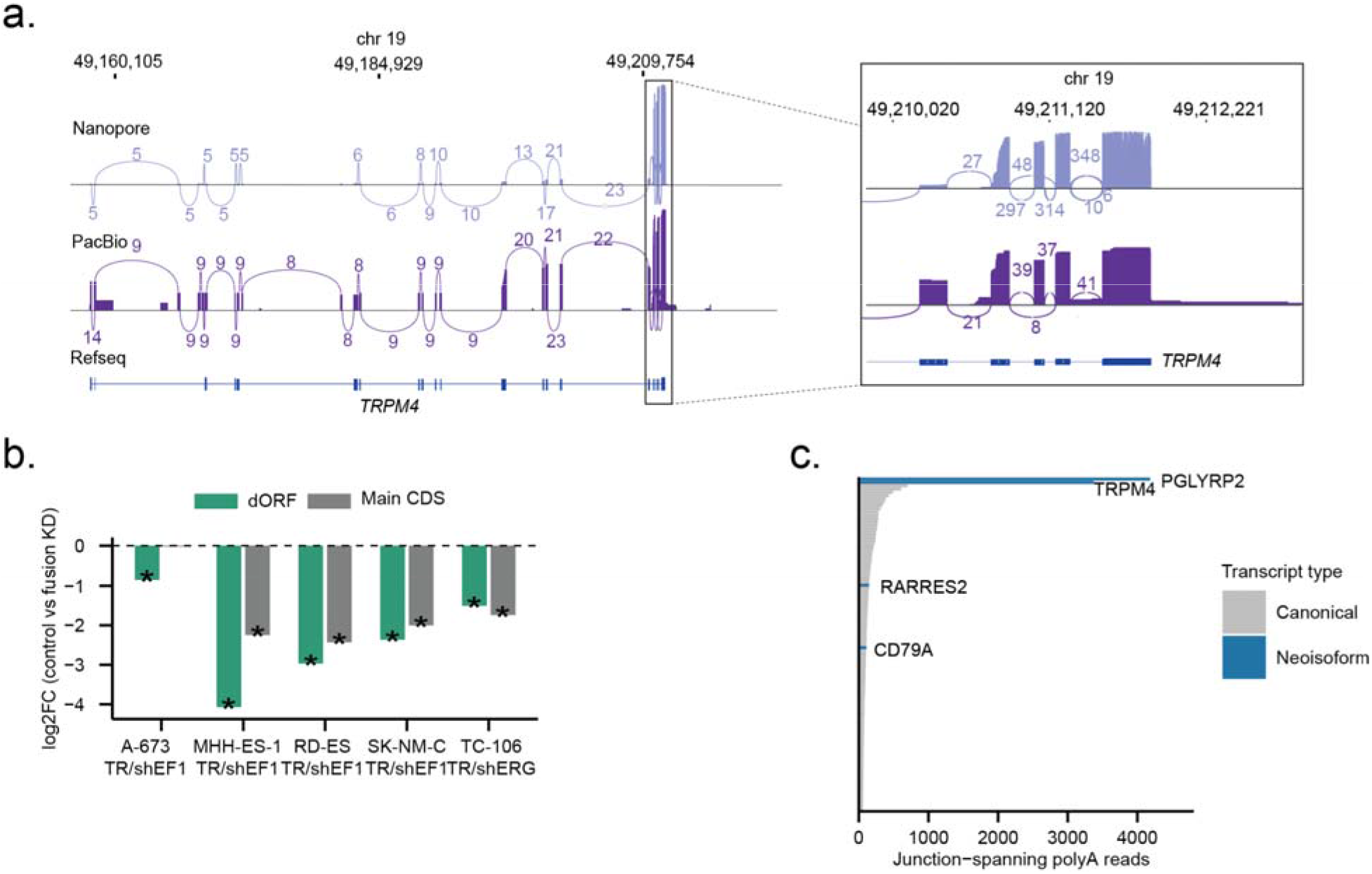
Supplementary information for *TRPM4-dORF* poly(A) readthrough RPF reads. **a.** Sashimi plot illustrating increased read coverage and splice junction usage within the terminal exons of *TRPM4* in Nanopore an PacBio long-read technologies. Junctions with fewer than 5 reads were not shown. **b.** Log fold-change in translational levels of the *TRPM4* main CDS and dORF following dox-treatment, shown for each individual cell line. **c.** Barplot indicating the number of BLAST-unique poly(A) RPF reads in EwS patients for the top 100 genes. Th highlighted genes are *EWSR1::ETS* regulated genes with neoisoforms.

**Figure S9.**
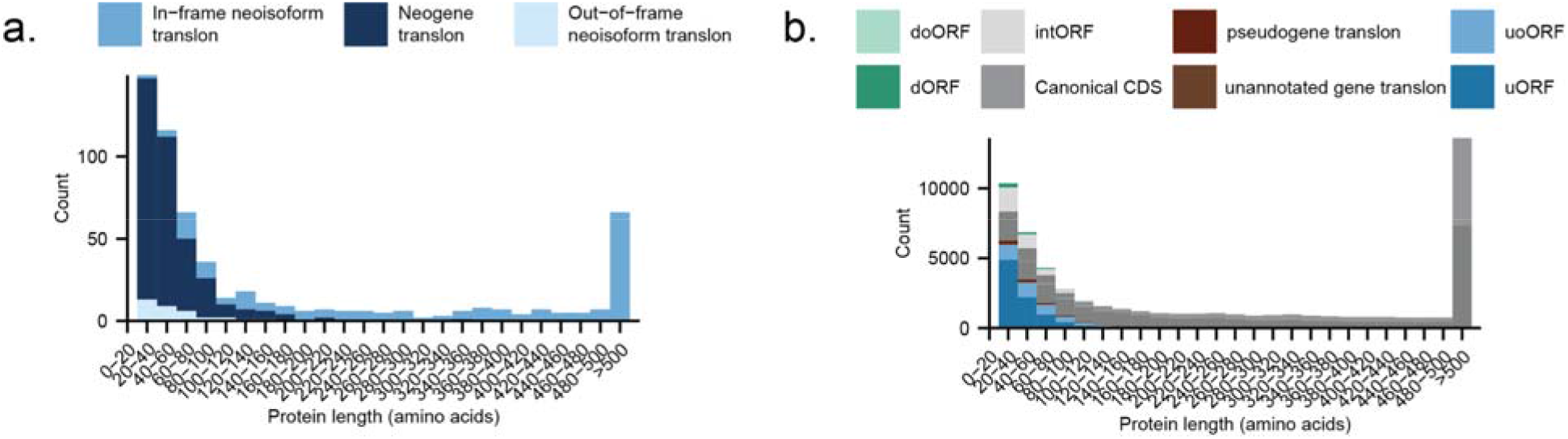
Amino Acid length of neoproteins and other nc-translon proteins. **a.** Histogram of neoprotein length, divided per translon category. **b.** Histogram of protein length of all non-canonical translon-encoded proteins.

**Figure S10.**
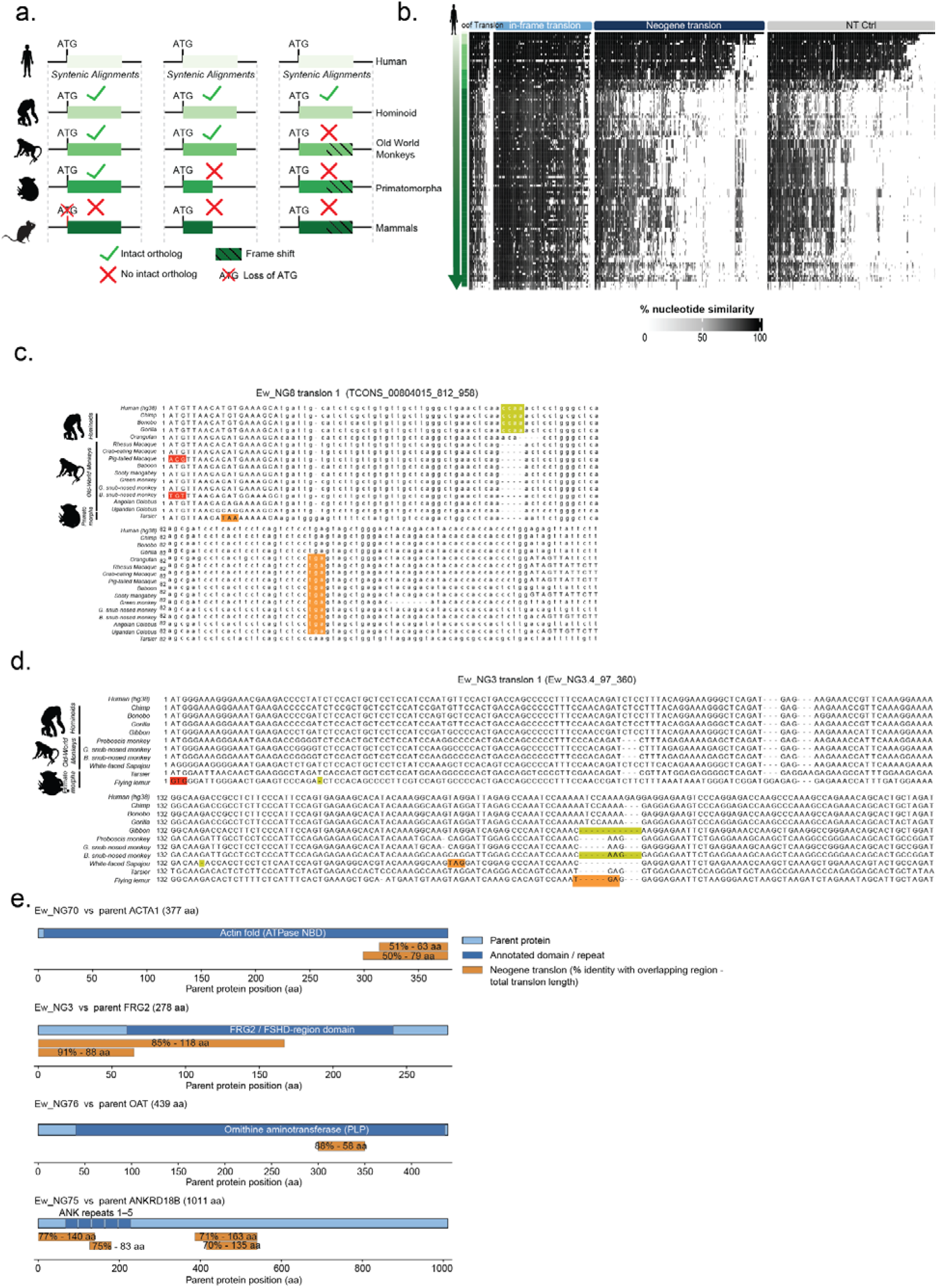
Supplementary information for evolutionary analysis. **a.** Schematic overview of the syntenic alignment approach used to assess translon conservation across species, illustrating criteria for intact orthologs: conserved ATG, maintained reading frame, and sequence length. **b.** Heatmap showing percentage of nucleotid similarity split by translon type. Each column is a translon, while each row is a pairwise-linked alignment in a species, ordered by phylogenetic age. **c-d**. Multiple alignments of Ew_NG8 translon 1 (TCONS_00804015_812_958) an Ew_NG3 translon 1 (Ew_NG3.4_97_360), ordered by phylogenetic age. Red nucleotide sequences indicate loss of ATG start codon, yellow indicate small insertions or deletions that change the reading frame and orange indicate premature stop codon. **e.** Schematic showing the 4 pseudogenic neogenes that contain translons with homology t their ancestral counterpart. Neogenes (in orange) show the overlapping region and the percentage of this overlapping region together with the length of the total translon including non-homologous regions. Percent identity is calculated over the aligned region between the neogene translon and its ancestral parent protein (local alignment); residues of the ORF outside this aligned region are not included.

**Figure S11.**
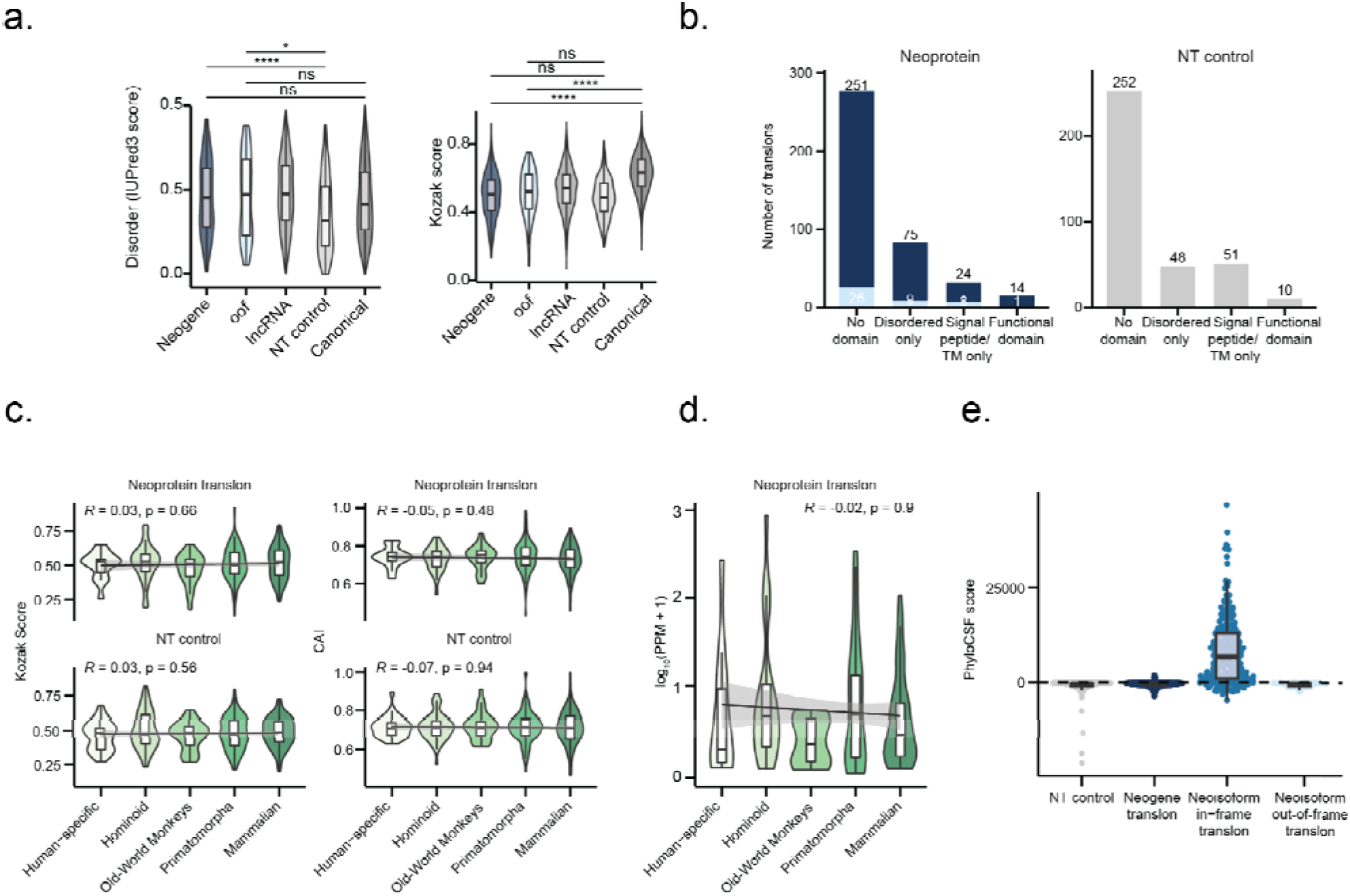
Biochemical features of neogenes and neoisoforms translons. a. Intrinsic disorder (IUPRED3 score) and Kozak scores for the (neo)proteins encoded by the translons (oof = Out-of-frame neoisoform) compared to canonical genes, lncRNA translons, length-matched non-translated controls (NT control). Significance was tested by the Wilcoxon Rank-Sum test (n.s., p > 0.05; *p ≤ 0.05; **p ≤ 0.01; ***p ≤ 0.001; ****p ≤ 0.0001). **b**. Number of Neoprotein (Neogenes and out-of-frame neoisoforms translons) with predicted protein domains as determined by InterProScan, classified as no domain, disordered only, signal peptide or transmembrane domain only, or recognizable functional domain. **c.** Kozak score and Codon Adaptation Index (CAI) across phylogenetic ages for Neoprotein translons and non-translated controls (NT control). Spearman correlation coefficient (*R*) and p-value are indicated. **d**. Translation levels (log (PPM + 1)) of neogene translons across phylogenetic ages. Spearma correlation (R) and p-value are indicated; gray shading represents the 95% confidence interval of the linear trend. **e**. PhyloCSF score for each translon type and the non-translated controls (NT control).

**Figure S12.**
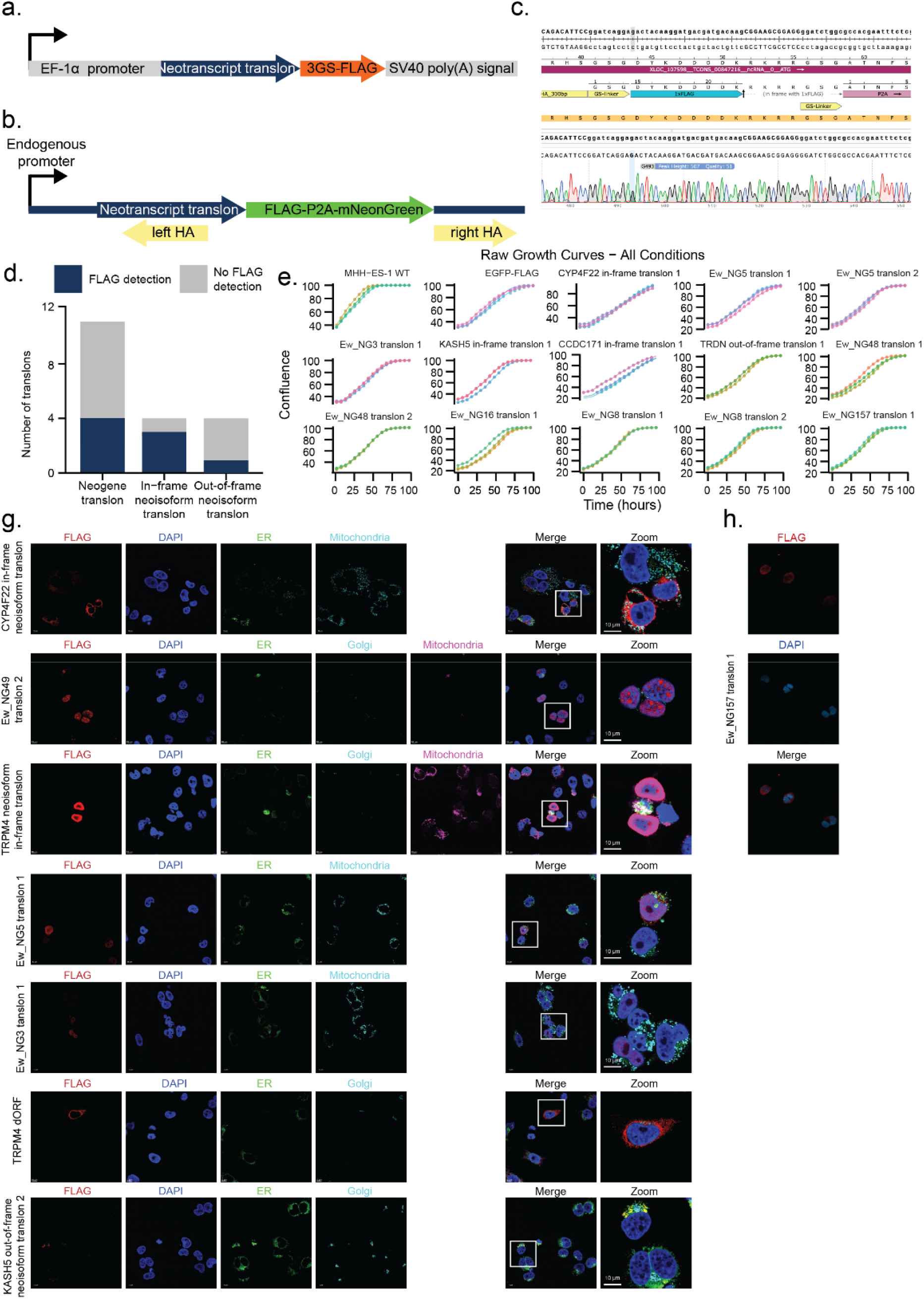
Transient overexpression and endogenous tagging. **a.** Schematic representation of the construct (pEF1α_T7_custom_MCS) used for transient overexpression in which the translon sequence was inserted. **b.** Schematic representation of the knock-in strategy. FLAG-P2A-mNeonGreen was inserted in-frame of Ew_NG5 translon under control of the endogenous promoter. Left and right arms (HA) flank the insert allowing for correct integration. **c.** Sanger sequencing results of gDNA of A-673 cells after endogenous tagging of the 3′ end of the neogene translon Ew_NG157. Sanger results show homozygous insertion of FLAG-tag without any frameshifts or mutations. **d.** Barplot showing the number of successfully transfected translon by translon type. **e.** Growth curves showing density of A-673 cells after transfection with transient overexpression plasmid containing the neogene and neoisoform translons. **f.** FLAG-Tag, DAPI, ER, Golgi and Mitochondria immunofluorescence stainings of all tagged neoproteins. **g.** FLAG-Tag, and DAPI immunostainings show nuclear localization of Ew_NG157 endogenously tagged neoprotein (XLOC_107598 TCONS_00847216_ncRNA_0_ATG).

**Figure S13.**
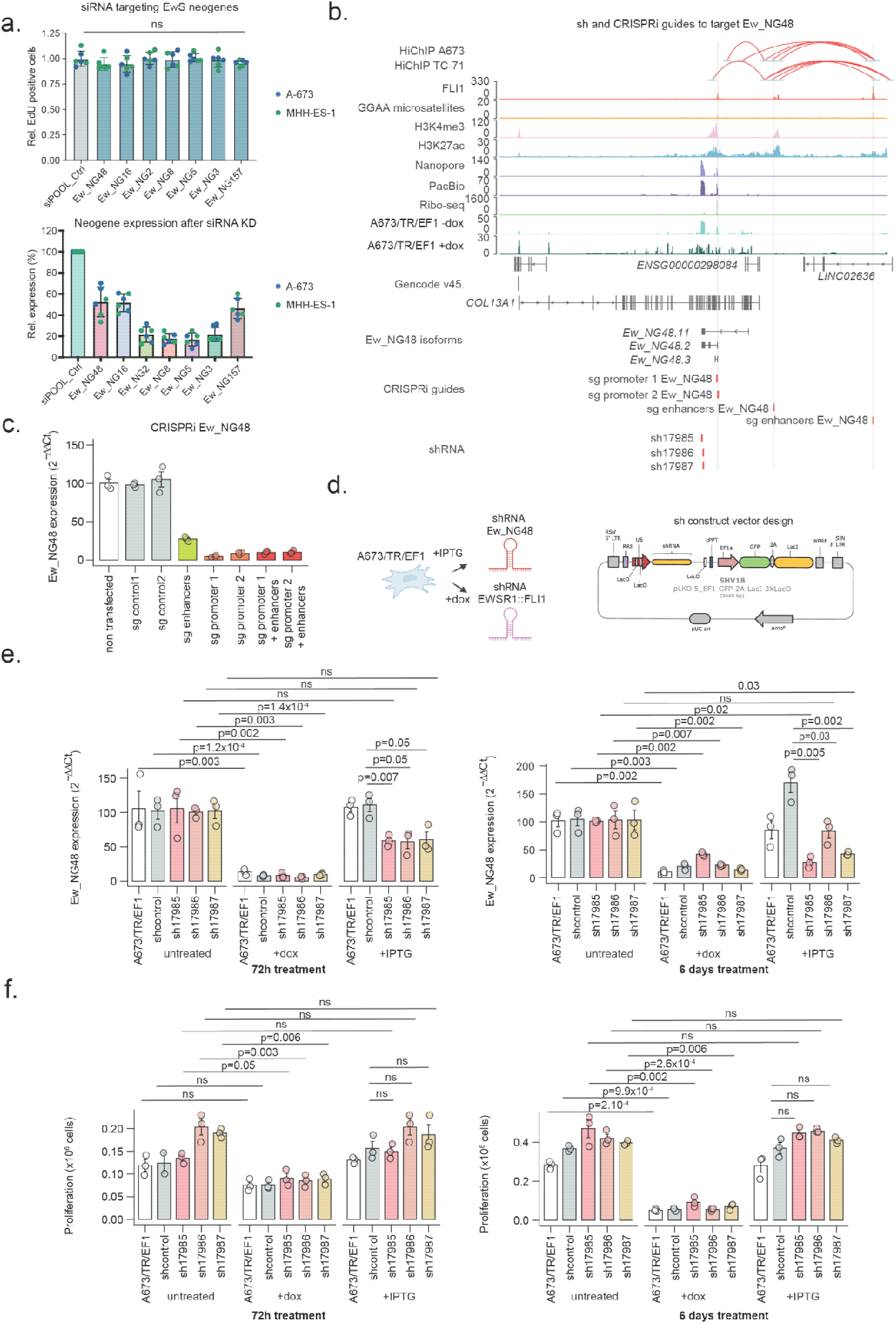
EwS neogenes knockdown (siRNA and shRNA) and CRISPR interference experiments. **a**. siRNA-mediated knockdown of 7 neogenes and control in A-673, and MHH-ES-1 EwS cell lines show no effect on EwS cells proliferation. Bar graphs below show the effect of siRNA-mediated knockdown on the expression level of the 7 neogenes. **b**. Genomic view of *Ew_NG48* in antisense orientation of *COL13A1* gene showing the design of shRNA guide and CRISPRi guides. CRISPRi guides target EF1-bound GGAA microsatellite at *Ew_NG48* TSS (sg promoter 1, sg promoter 2) or within enhancer-promoter chains linked to *Ew_NG48* (sg enhancers). **c.** CRISPR interference targeting EF1-bound GGAA microsatellites shows *Ew_NG48* downregulation. Both guides targeting EF1-bound GGAA microsatellite at TSS or within promoter-enhancer chains result in Ew_NG48 downregulation. Experiment was done only once, triplicates of qPCR are displayed. **d**. Schematic of the plasmid construct containing IPTG-inducible shRNAs targeting the *Ew_NG48* transcript. A673/TR/shEF1 harbor a dual-inducible shRNA system in which dox induces an shRNA targeting EF1 fusion protein^110^ while IPTG induces expression of an shRNA targeting *Ew_NG48*. **e**. *Ew_NG48* Expression levels (2^-ΔΔCt^) in A673/TR/shEF1 cells after 72h or 6 days of dox or IPTG treatment compared with untreated cells. Experiments were performed in biological triplicate with three qPCR technical replicates each. Statistical comparisons were performed using Student’s *t*-test. **f**. A673/TR/shEF1 cell counts after 72h or 6 days of dox or IPTG treatments compared with untreated cells. Experiments were performed in biological triplicate with three technical replicates each. Statistical comparisons were performed using Student’s *t*-test.

